# DNA-origami line-actants control domain organisation and fission in synthetic membranes

**DOI:** 10.1101/2023.01.09.523307

**Authors:** Roger Rubio-Sánchez, Bortolo Matteo Mognetti, Pietro Cicuta, Lorenzo Di Michele

## Abstract

Cells can precisely program the shape and lateral organisation of their membranes using protein machinery. Aiming to replicate a comparable degree of control, here we introduce DNA-Origami Line-Actants (DOLAs) as synthetic analogues of membrane-sculpting proteins. DOLAs are designed to selectively accumulate at the line-interface between co-existing domains in phase-separated lipid membranes, modulating the tendency of the domains to coalesce. With experiments and coarse-grained simulations, we demonstrate that DOLAs can reversibly stabilise two-dimensional analogues of Pickering emulsions on synthetic giant liposomes, enabling dynamic programming of membrane lateral organisation. The control afforded over membrane structure by DOLAs extends to three-dimensional morphology, as exemplified by a proof-of-concept synthetic pathway leading to vesicle fission. With DOLAs we lay the foundations for mimicking, in synthetic systems, some of the critical membrane-hosted functionalities of biological cells, including signalling, trafficking, sensing, and division.

## Introduction

Biological membranes coordinate numerous pathways critical to life, from trafficking to signal transduction and cellular motility,^1^ many of which rely on regulating the distribution and interactions of membrane machinery. Among other regulatory principles, cells are believed to exploit membrane phase separation and critical composition fluctuations to dynamically generate local heterogeneity.^2–10^ Cell membranes are also known to contain inclusions that can accumulate at the line-interface between co-existing lipid domains,^11–15^ and relax the associated line tension.^16–18^ It has been proposed that cells use these *line-actants* as an additional means to dynamically control membrane phase behaviour and the lateral organisation of membrane inclusions^19^ underpinning, for instance, the function of signalling hubs^20,21^ and the generation of endo/exosomes. ^22,23^

Artificial cell science aims to construct, from the bottom-up, synthetic analogues of biological cells that mimic their sophisticated behaviours,^24,25^ and are expected to widely impact next generation diagnostics, therapeutics and bioprocessing.^26–28^ Often constructed from lipid bilayers, ^29^ synthetic-cell membranes have been engineered to replicate an array of biomimetic responses,^30^ including mechanotransduction,^31^ energy conversion, ^32–34^ and membrane deformation.^35^

In parallel to solutions relying on reconstituted membrane proteins,^32–34^ fully synthetic DNA nano-devices,^36^ anchored to the bilayers by means of hydrophobic tags, have emerged as a versatile toolkit for bio-membrane engineering,^37–39^ having enabled the design of biomimetic pathways for transport,^40–43^ trafficking,^44,45^ cell adhesion, ^46,47^ tissue formation,^48–50^ signal detection,^51^ membrane remodelling, ^52–55^ and surface patterning. ^56–59^

Despite these advances, the design of pathways enabling systematic control over the local structure and composition of synthetic-cell membranes, to the same degree of what is afforded by extant biological machinery, remains an elusive task.

Here we introduce DNA-Origami Line-Actants (DOLAs) to control the formation, stability and three-dimensional morphology of lipid domains on synthetic bilayers. Leveraging the modularity of amphiphilic DNA nanotechnology,^56^ we designed DOLAs to selectively accumulate at the line-interface between co-existing domains on Giant Unilamellar Vesicles (GUVs). Using a combination of experiments and coarse-grained simulations, we show that, similar to interfacial inclusions in three dimensions, DOLAs modulate the tendency of lipid domains to coalesce, stabilising two-dimensional analogues of Pickering emulsions. The devices can be de-activated upon exposure to a molecular stimulus, triggering domain coarsening and unlocking the sought dynamic control over lateral membrane organisation. Combined with osmotic unbalance, we demonstrate DOLAs can form the basis of a synthetic pathway leading to membrane budding-off and fission, exemplifying three-dimensional morphological control.

Owing to their modular design and robust working principle, we argue that DOLAs could constitute a versatile toolkit for synthetic-cell membrane engineering, allowing us to take full advantage of the rich phenomenology of lipid phase separation and design ever-more advanced membrane-hosted functionalities.

## Results and Discussion

### Engineering line-actants with DNA nanostructures

Our design of choice for DNA line-actants, depicted schematically in Fig. 1a, exploits the versatility of the Rothemund Rectangular Origami (RRO) as a “molecular breadboard”, with an array of regularly spaced binding sites. ^61–64^ The rectangular tiles (Supplementary Fig. 1 and Supplementary Table 1) feature two sets of either 6, 12 or 24 single-stranded (ss) DNA overhangs extended from the same face of the origami, as shown in Fig. 1b. The sticky ends, with sequence *δ*1* or *δ*2*, can respectively bind complementary overhangs on double-stranded (ds) DNA *anchoring modules* functionalised with two cholesterol moieties (dC) or a single tocopherol (sT). ^56^ We have recently determined that dC and sT modules display, respectively, thermodynamic preferences for accumulating within liquid-ordered (*L*_o_) and liquid-disordered (*L*_d_) domains of phase-separated membranes. Specifically, transporting a dC module from *L*_d_ to *L*_o_ produces a moder-ately favourable free-energy shift 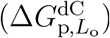 of ∼ − 0.8 *k*_B_*T*, while sT has a stronger affinity for disordered domains 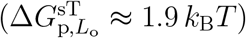.^56^ The two sets of dC and sT modules were distributed on opposite halves of the tiles, with spacing of *l*∼ 25 − 30 nm between the two distinct sets of anchors (Fig. 1b). Because *l* is comfortably greater than the estimated width of the boundary separating *L*_o_ and *L*_d_ phases, (*ξ* ∼ 8 nm, Supplementary Note I), free-energy minimisation is expected to drive the accumulation of the tiles across line-interfaces, accommodating each set of anchors in their respective preferred phase (Fig. 1a). We produced three DOLA designs, with either 6*×*, 12*×* or 24*×* anchors of each type, expecting an increase in the free-energy gain associated to line-accumulation for larger number of anchors (Fig. 1c).

**Figure 1:**
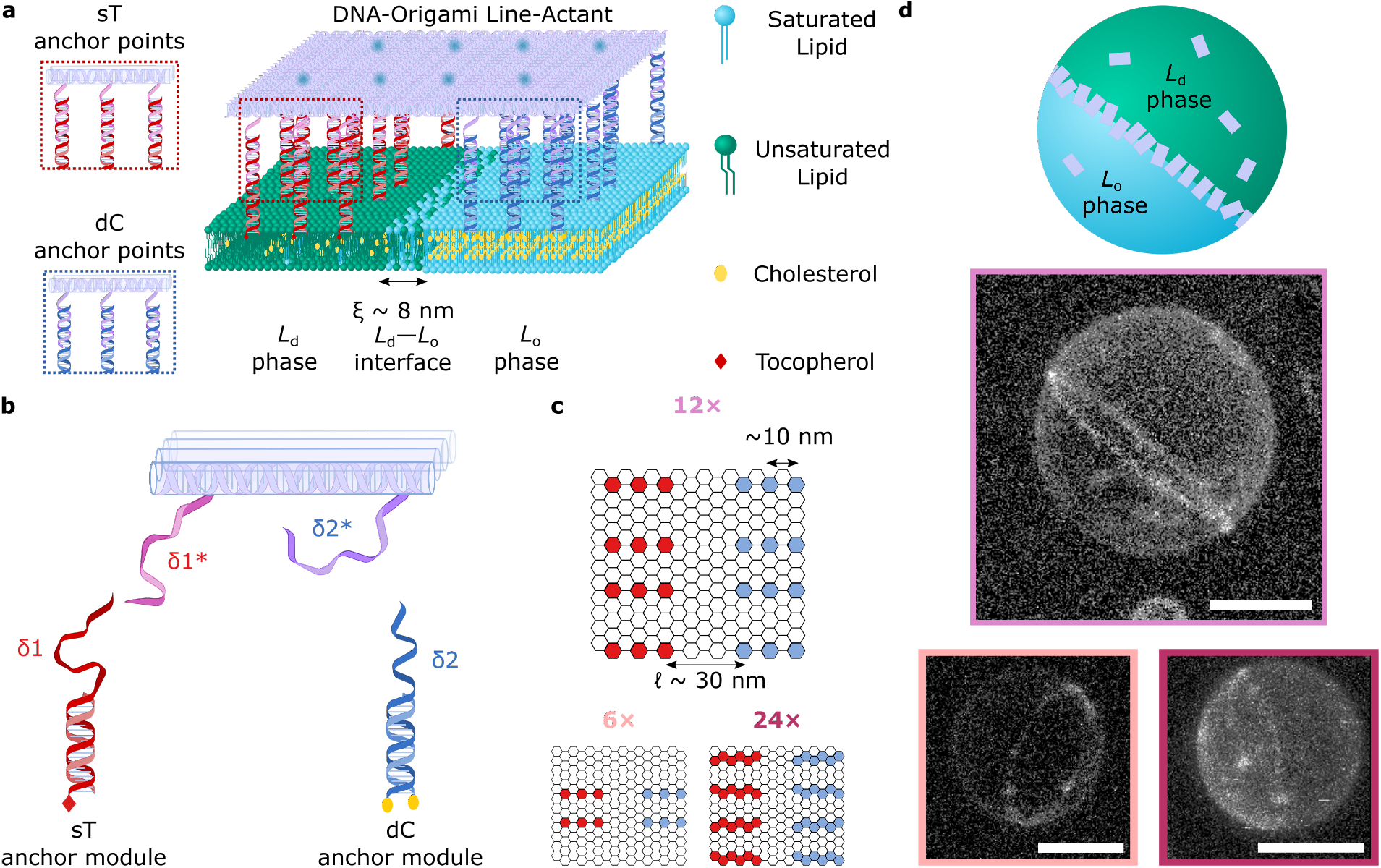
DNA-Origami line-active agents enrich the line-interface of phase-separated GUVs. **a** Schematic representation of a multi-component lipid bilayer membrane composed of saturated and unsaturated lipids mixed with sterols, displaying liquid-ordered (*L*_o_) and liquid-disordered (*L*_d_) phase co-existence. The outer leaflet of the membrane is decorated with a DNA-Origami Line-Actant (DOLA), where amphiphilic anchor-modules bearing single-tocopherol (sT) and double-cholesterol (dC) motifs enrich their preferred (*L*_d_ and *L*_o_, respectively) phases, driving accumulation of the DOLA at the *L*_d_ − *L*_o_ interface. **b** Selected staples on the origami (see Supplementary Table 2) were extended to include overhang domains *δ*1* and *δ*2*, with complementarity to dC (blue) or sT (red) anchor modules, respectively (Supplementary Tables 3 and 4). Note that tiles are expected to have all of their *δ*_1_ and *δ*_2_ overhangs saturated with sT and dC anchoring modules, owing to the strong affinity between overhangs (see Supplementary Note III). We also expect partially bound states, in which tiles are linked to the membrane through a sub-set of the available anchors, to be thermodynamically unlikely. **c** Hexagonal grids depict the arrangement of staples across the origami plates, where each hexagon corresponds to the 3’ terminus of a staple. We positioned sets of 6× (light pink), 12× (pink), or 24× (magenta) overhangs targeting the same type of anchor-points. The two sets of anchors are separated by a distance *l* ∼ 30 nm (6× and 12×) or *l* ∼ 25 nm (24×). **d** (Top) Schematic representation a de-mixed GUV where the *L*_d_ − *L*_o_ interface is enriched with DOLAs. (Bottom) Representative 3D views of phase-separated vesicles with a Janus-like morphology showing line-accumulation of fluorescent (Alexa488) 6× (light pink), 12× (pink), or 24× (magenta) DOLAs, as reconstructed from a confocal z-stack using Volume Viewer (FIJI ^60^) with contrast enhancement (see Supplementary Fig. 3 for reconstructions without contrast enhancement and confocal equatorial cross sections highlighting line accumulation). Scale bars = 10 *μ*m.

DOLAs were further labelled with fluorescent (Alexa488) beacons, located on the face oppo-site to that hosting the anchors (Supplementary Fig. 2 and Supplementary Table 5), thus allowing us to monitor their distribution by means of confocal microscropy. Accumulation of the three DOLA designs at the *L*_d_–*L*_o_ line-interface in phase-separated GUVs (DOPC/DPPC/Chol 2:2:1) is demonstrated in Fig. 1d with confocal 3D reconstructions, where the equatorial boundary separating the two hemispherical domains is clearly delineated by the fluorescent plates.

A custom-built image segmentation routine, detailed in Supplementary Note II and Supplementary Fig. 4, was applied to confocal data to sample the DOLAs fluorescence-intensity profile across line-interfaces, as sketched in Fig. 2a. The resulting curves, shown in Fig. 2b, display clear peaks at the *L*_d_–*L*_o_ boundary location (*x* = 0), confirming line-accumulation. As detailed in Supplementary Note II, fitting of the diffraction-limited fluorescent peak and base-line signals from the surrounding *L*_d_ and *L*_o_ regions allowed us to estimate a line-partitioning coefficient *K*_p,int_, defined as the ratio between the surface density of origami at the line-interface and the average origami surface density on the entire GUV.

**Figure 2:**
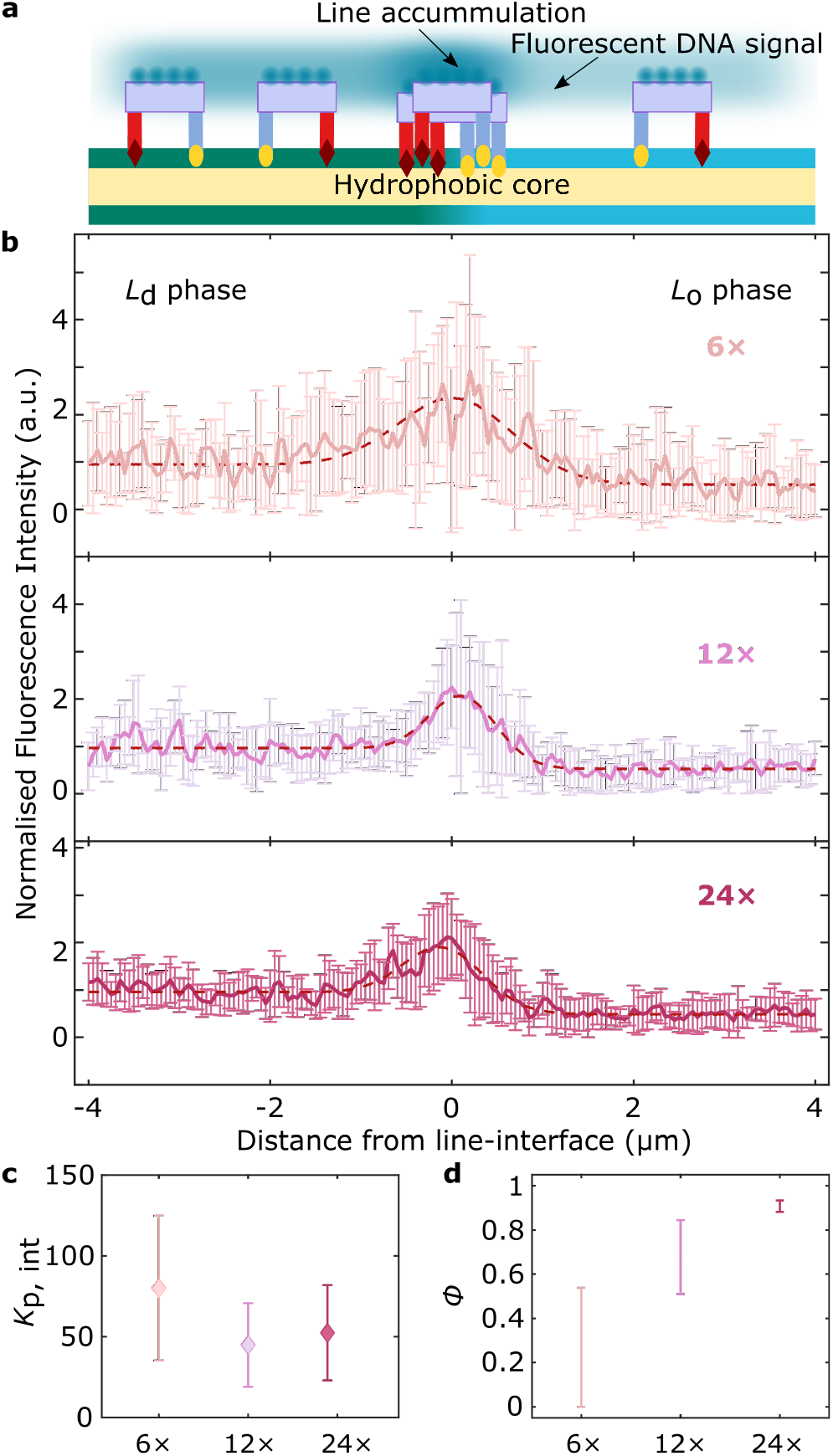
Line-accumulation of DOLAs is regulated by the number of anchors. **a** Schematic depiction of a de-mixed lipid membrane decorated with DOLAs, where fluorescent Alexa488 beacons enable the quantification of origami distribution from fluorescence intensity profiles (Supplementary Note II). **b** Line-interface accumulation conveyed as average fluorescence intensity *±* standard deviation for line interfaces decorated with 6×, 12×, or 24× DOLAs (sampled interfaces *n* = 12, 10 and 10, respectively). The profiles, as a function of distance from the line-interface (*x* = 0), have been normalised to the mean *L*_d_-phase intensity. Red dashed lines are fits to Eq. S2 (Supplementary Note II), which models the experimental, diffraction-blurred, fluorescence intensity profile. **c** Line-partitioning coefficients (*K*_p,int_) of DOLAs featuring sets of either 6×, 12×, or 24× dC/sT anchor-points. **d** Fraction of line-interface occupied (*ϕ*) by 6×, 12×, or 24× line-actants, computed using the experimental *K*_p,int_ in our numerical line-adsorption model (see Supplementary Note III and Supplementary Figs. 5, 6, 7, and 8.)

Figure 2c summarises the experimentally determined *K*_p,int_ for the three tested designs, rendering similar results for the 12× and 24× tiles, with a slightly larger value for the 6× variant. Line-interface adsorption models, outlined in Supplementary Note III (see associated Supplementary Fig. 5 and Supplementary Tables 6, 7, and 8), enable the estimation of *K*_p,int_ for the three DOLA designs. We fitted the theoretical estimates to experimentally-determined *K*_p,int_ values, using the overall surface coverage of the origami on the GUVs, *σ*, as fitting parameter.

The obtained estimates of 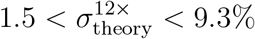 and 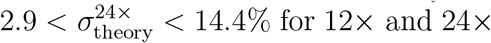 for 12× and 24×, respectively, are in good agreement with the nominal experimental surface coverage *σ*_exp_ ∼ 5% (Supplementary Fig. 6). For the 6× design, we estimated 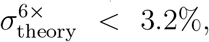, slightly below the nominal experimental value and reflecting the larger observed *K*_p,int_ and the smaller absolute fluorescence-intensity values (see Supplementary Note III and Supplementary Fig. 7). A smaller than expected *σ* is likely the result of a decreased membrane-affinity, consistent with the smaller number anchor-points available for the 6× tiles. It should further be noted that RRO tiles are prone to bending in the direction orthogonal to that of the double helices,^65^ possibly leading to thermal fluctuations in curvature while tiles are in the bulk. These fluctuations would be suppressed upon confining the tiles to the membrane, which would carry an entropic penalty to tile-membrane binding. It is natural to expect that anchoring overhangs may limit the amplitude of shape fluctuations in the bulk, implying that designs with a higher number of anchoring points, *i*.*e*. 24× and 12×, would be less affected by the confinement entropic penalty compared to the (arguably) more flexible 6× design. Possible differences in spontaneous (equilibrium) curvature of the tile designs^66^ may also impact membrane-tile affinity. Fitted line-adsorption models allowed us to estimate the fraction of the line-interface occupied by DOLAs, *ϕ* (defined in Supplementary Note III; see Supplementary Fig. 5), summarised in Fig. 2d. Expectedly, we note an increase in *ϕ* with increasing number of anchors, with the strongly-binding 24× approaching saturation and the 6× leaving much of the line unoccupied, hence demonstrating control over the degree of line-accumulation by design. Note that the uncertainties in the estimate of the line-interface width *ξ* propagate to the values obtained for *K*_p,int_ (see Supplementary Note III). As demonstrated in Supplementary Fig. 8, however, relative trends with respect to tile design are robust, while changes to the values of *ϕ* are negligible.

### DOLAs stabilise two-dimensional Pickering emulsions

Having demonstrated that DOLAs can programmably accumulate at the boundary between co-existing lipid phases, we sought to verify whether accumulation leads to stabilisation of the line-interface, as hypothesised for biological line-actants.^14^ To this end, we decorated the phase-separating GUVs with 12× DOLAs at a nominal surface coverage of ∼ 30%, further promoting line-interface saturation (*ϕ* ∼ 0.75−0.87). The GUVs, fluorescently labelled with *L*_d_-partitioning TexasRed-DHPE for ease of visualisation, were heated to ∼ 37^°^C, well above their miscibility transition temperature of *T*_m_ ∼ 33 *±* 1^°^C,^68^ to induce lipid mixing. After incubating at high temperature for ∼ 5 minutes, the sample was quenched back to room temperature (*T* = 25^°^C), leading to the nucleation of *L*_o_ or *L*_d_ domains in a background of the opposite phase.

In non-functionalised GUVs, domains rapidly coarsen until most vesicles display two quasi-hemispherical domains, resulting in Janus-like morphologies (Fig. 3a, bottom). However, in DOLA-functionailsed GUVs, coalescence was arrested leading to morphologies with a large number of small, stable domains, analogous to two-dimensional Pickering emulsions (Fig. 3a, top). To quantify the domain-stabilising ability of DOLAs, we analysed epifluorescence micrographs collected ∼ 3 hrs after quenching, and extracted the fraction *F* of non-Janus vesicles, namely those that failed to relax into the two-domain morphology. Results, summarised in Fig. 3b, clearly demonstrate how most line-actant decorated GUVs display Pickering morphologies (Fig. 3a, top). In turn, the Janus morphology dominates when GUVs lack either dC or sT anchor modules, no tiles are included, or if the origami are prepared omitting the overhangs to target either dC or sT. The latter controls confirm that domain stabilisation emerges solely thanks to line-interface accumulation of the rationally designed DOLAs, rather than as a result of non-specific effects associated to the individual membrane inclusions.

**Figure 3:**
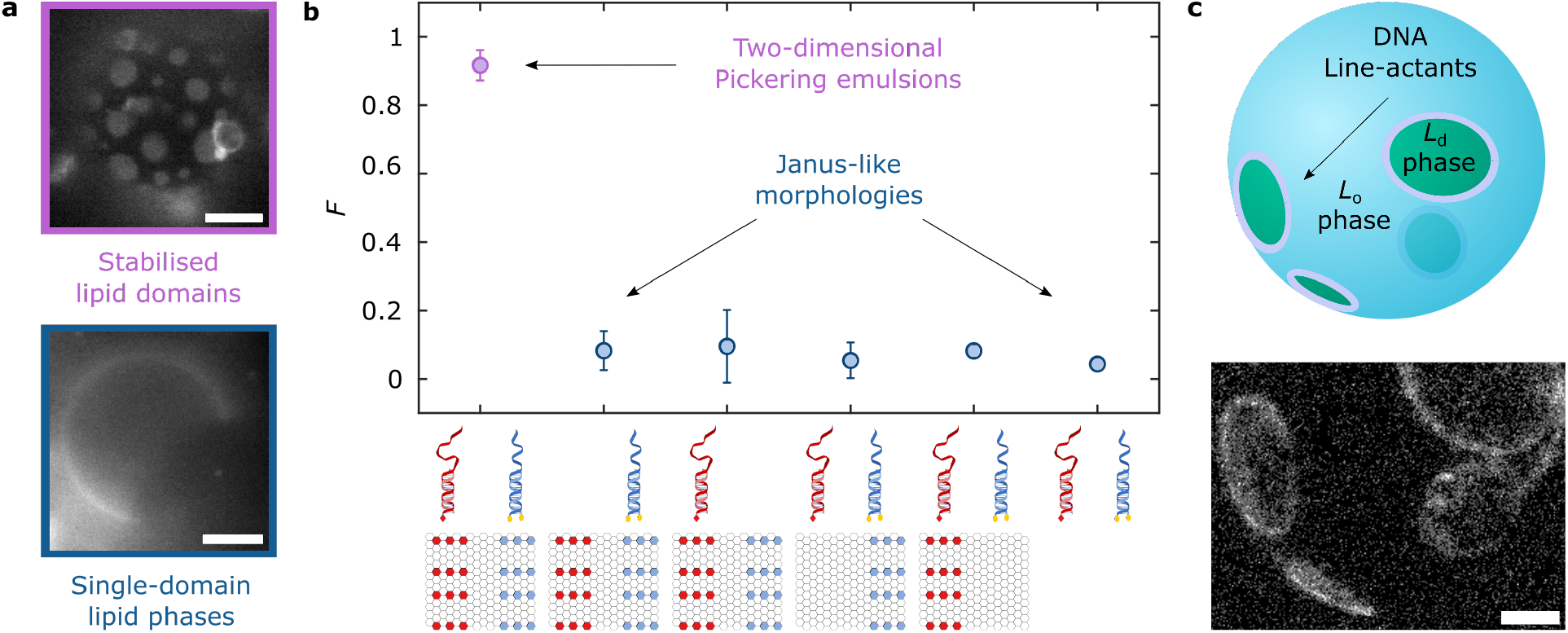
DOLAs stabilise two-dimensional analogues of Pickering emulsions. **a** Representative epifluorescence micrographs of de-mixed GUVs showing: (top) stable lipid domains ≈ 3 hours after phase separation in the presence of 12× DOLAs; (bottom) GUV lacking line-actants that equilibrated to produce a Janus morphology with two quasi-hemispherical *L*_d_ and *L*_o_ domains. Fluorescent marker is Texas Red-DHPE, which partitions to the *L*_d_ phase. **b** Programmable, domain-stabilising, activity of DOLAs as conveyed by the fraction, *F*, of GUVs exhibiting more than two domains with respect to that of Janus vesicles. As shown in the graphical legend, besides the complete DOLAs (*n* = 363 GUVs), we exploited the modularity of the platform to produce various control functionalisation schemes expected to lack line-active behaviour, including omitting either sT (*n* = 97 GUVs) or dC (*n* = 187 GUVs) modules, using tiles that lack overhangs targeting sT (*n* = 125 GUVs) or dC (*n* = 165 GUVs), and omitting tiles altogether (*n* = 160 GUVs). **c** Line-accumulation of DOLAs scaffolds lipid domains and confers them with stability against coalescence, as shown with a 3D view of a reconstructed vesicle after heating above and quenching below the miscibility transition temperature. Interfacial accumulation of fluorescent (Alexa488) 12× DOLAs readily shows stable domains rendered from a representative confocal z-stack using Volume Viewer (FIJI ^60^) with contrast enhancement (see Supplementary Fig. 9 for reconstructions without contrast enhancement). All scale bars = 10 *μ*m.

As further confirmation that domain stabilisation is underpinned by interfacial accumulation of the origami, vesicles lacking TexasRed-lipids and decorated with fluorescent DOLAs were subjected to an analogous heating-cooling program and inspected with confocal microscopy. Besides detecting domain stabilisation, line-accumulation was confirmed at the boundaries of the stabilised lipid domains, as demonstrated in Fig. 3c with a 3D confocal reconstruction.

### Coarse grained simulations capture DOLA line-accumulation and domain stabilisation

To further validate our line-actant engineering strategy, we conducted Monte Carlo (MC) simulations using a coarse-grained representation of the experimental system. As presented in Fig. 4a, and detailed in Supplementary Note IV, we defined a two-dimensional Ising model on a triangular lattice, which displays phase co-existence at sufficiently low temperature. We interpret the phase rich in spin *s* =−1 (green) as *L*_d_, and that rich in *s* = 1 (blue) as *L*_o_. Rectangular tiles were included with anchor-points arranged on the lattice according to their nominal position on the DOLAs. Interaction free-energies between dC/sT anchors and their lipid micro-environment were modelled in the system’s Hamiltonian as 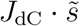 and 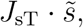, respectively, where 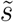 is the sum of the spins over the lattice site hosting the anchor and its six nearest neighbours 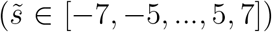. The anchor-coupling constant values were chosen as *J*_sT_ = 0.136 *k*_B_*T* and *J*_dC_ = 0.057 *k*_B_*T*, reflecting the experimentally-determined partitioning free-energies of the modules (see Supplementary Note IV and ref.^56^).

**Figure 4:**
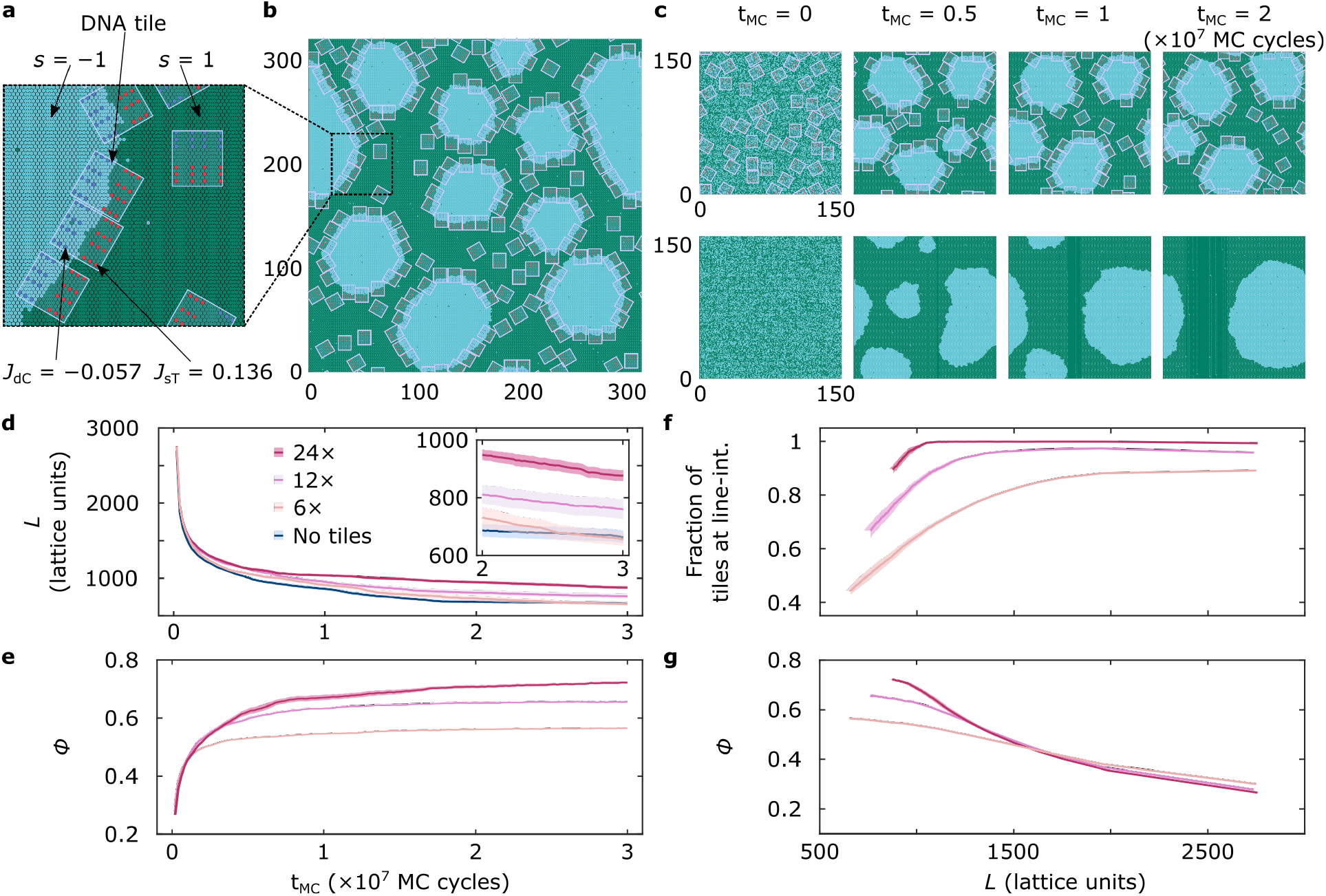
Coarse-grained simulations replicate DOLA line-accumulation and domain stabilisation. **a** Representation of the developed Ising model with hexagonal unit cells carrying spins of *s* = 1 (green) and *s* = −1 (blue). Neighbouring spins interact through a coupling parameter *J* = 0.55 *k*_B_*T*, chosen to roughly replicate the expected experimental line tension ∼ 1 pN. ^67^ Anchor-points on the tiles are arranged based on the 12× DOLA design. Each anchor is located at an Ising lattice point and contributes with a free-energy term 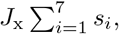, where *s*_*i*_ indicates the spins of the lattice point hosting the anchor and the 6 nearest neighbors, and *J*_x_ is a coupling constant taking different values for sT (*J*_sT_ = 0.136 *k*_B_*T*) and dC (*J*_dC_ = − 0.057 *k*_B_*T*) to account for the different partitioning tendencies. **b** Representative snapshot of a simulation trajectory after *t*_MC_ = 2 (× 10^7^ MC cycles), where tiles stabilise blue domains by accumulating at their boundary. Box size: 320 × 372 lattice points. **c** Snapshots exemplifying the time-evolution of representative simulations, one including 12× DOLAs, where multiple domains retain stability even after *t*_MC_ = 2 × 10^7^ MC cycles (top), and the second lacking tiles and equilibrating to form a single domain (bottom). Box size: 160 × 172 lattice points. **d** Time dependence of the overall length of the line-interface (*L*) in simulations containing the three tested DOLA designs or no tiles, quantifying DOLA-induced interface stabilisation. **e** Line-accumulation of 6×, 12×, and 24× DOLAs conveyed by the time-evolution of fraction of line-interface (*ϕ*) occupied by tiles. A tile is said to be at the line when featuring at least 80% of favourable anchor-spin interactions relative to the total contacts (Supplementary Fig. 11). **f** Fraction of the tiles present in the system located at the interface, showing that designs remain pinned at the line more readily as the latter shrinks. **g** Fraction of the line-interface (*ϕ*) occupied by tiles as a function of line length (*L*). The monotonic decrease is sharper for stronger-binding tiles. Panels **d**–**g** show mean (solid line) *±* standard error on the mean (shaded regions) computed from *n* = 15 independent simulation trajectories. When errors are not visible, their values are smaller than the thickness of the solid line.

To replicate the experimental protocol, we prepared high-temperature systems with uniform spin (and tile) distributions, before quenching to a lower temperature at which demixing occurs. The quenching temperature was chosen to replicate the experimental line tension of ∼ 1 pN^67^ (see Supplementary Note IV). The system was simulated with Kawasaki dynamics,^69^ conserving the number of *s* = − 1 and *s* = 1 spins, initially set to a ratio of 2/3. The tiles, present at a surface coverage *σ* = 30%, readily localised at the boundaries of the blue domains emerging after quenching, leading to their stabilisation, (see Fig. 4b and Supplementary Movie 1). Figure 4c compares the time-evolution of systems featuring and lacking 12× tiles. When tiles were present, coarsening was slowed down and small domains persisted at long simulation times (*t*_MC_ = 2 × 10^7^ steps). Conversely, over the same time window, the system lacking the tiles achieved complete coarsening, resulting in a morphology with one domain of each phase, analogous to the Janus GUVs (Supplementary Movies 2 and 3, re-spectively). Supplementary Fig. 10 compares frequency histograms of the number of domains in 15 runs at various *t*_MC_, confirming the ability of the tiles to slow down domain coarsening, in line with experimental findings outlined in Fig. 3.

Despite its coarse-grained nature, simulations offer insights on the different mechanisms lead-ing to domain coarsening, and on how these are influenced by the presence of DOLAs. Generally, we observed coarsening followed two distinct pathways: domain coalescence and Ostwald ripening. ^70^ Simulations suggest that the presence of tiles slows down the former mechanism, by offering a steric barrier that prevents the interfaces of two distinct domains from com-ing in sufficient proximity and undergo coalescence. Instead, Ostwald ripening, supported by the exchange of particles (spins) between domains *via* diffusion through the background phase, occurs regardless of the presence of tiles. This second coarsening process is solely responsible for the decrease in total line-length observed in the presence of tiles at very long simulation times (Fig. 4d, inset), when no further domain coalescence is observed. In experiments, tile-decorated GUVs display stable 2D Pickering emulsions ∼ 3 hours after quenching, when most non-functinonalised vesicles have fully equilibrated (Fig. 3b), hinting that domain coalescence may be the primary mechanism leading to coarsening, with Ostwald ripening playing a comparatively minor role.

Figure 4d compares the simulated time-evolution of the overall line-interface length (*L*) for systems featuring each DOLA design with those lacking tiles. At late simulation times, the interface length is higher for 24×, followed by 12× and 6×, with the latter design showing values identical to the tile-less control, confirming the expected link between number of anchors and line-stabilising ability. See also Supplementary Fig. 11 and the associated Supplementary Movies 4 and 5 for representative simulation runs with 24× and 6× DOLAs.

Consistently, as shown in Fig. 4e, the simulated tile-occupied fraction of the line (*ϕ*) converges to values that monotonically increase with the number of anchors, in agreement with the experimental/theoretical findings outlined in Fig. 2d. Analogous trends are noted when monitoring the time-dependent fraction of tiles at the line-interface, found to increase with anchor-point number (Supplementary Fig. 13). Further insights can be gathered from Fig. 4f, where we plot the fraction of tiles at the line as a function of interface length. We note that, as *L* decreases due to domain coarsening, strongly binding 24× tiles remain more persistently pinned at the interface, while 6× tiles are readily expelled. Finally, in Fig. 4g, we explicitly explore the correlation between *ϕ* and line length. As *L* shrinks, a plateau in *ϕ* Isapproached for 6× tiles able to readily desorb from the line, while a correlation is retained for strongly-pinned 24× DOLAs.

### Fueling lipid domain re-organisation with dynamic DNA line-actants

The ability of DOLAs to regulate domain coarsening can be readily coupled to nanostructure re-configurability afforded by toeholding reactions,^71–73^ thus enabling dynamic control over membrane lateral organisation. To demonstrate this functionality, 12× line-actants were modified with a 6-nt toehold domain of sequence *α* on the 3’-end of cholesterol-targeting overhangs (see Supplementary Table 9). As depicted schematically in Fig. 5a, a chemical Fuel, in the form of an oligonucleotide with sequence *α*\**δ*2, can selectively displace the dC-anchoring modules from the tile through a toe-holding reaction (see Supplementary Fig. 14 for an agarose gel confirming the sought molecular response). When removing the dC anchors, DOLAs are expected to lose their line-interface affinity and instead partition to *L*_d_ driven by the remaining sT moieties, negating the ability of the tiles to stabilise lipid domains.

**Figure 5:**
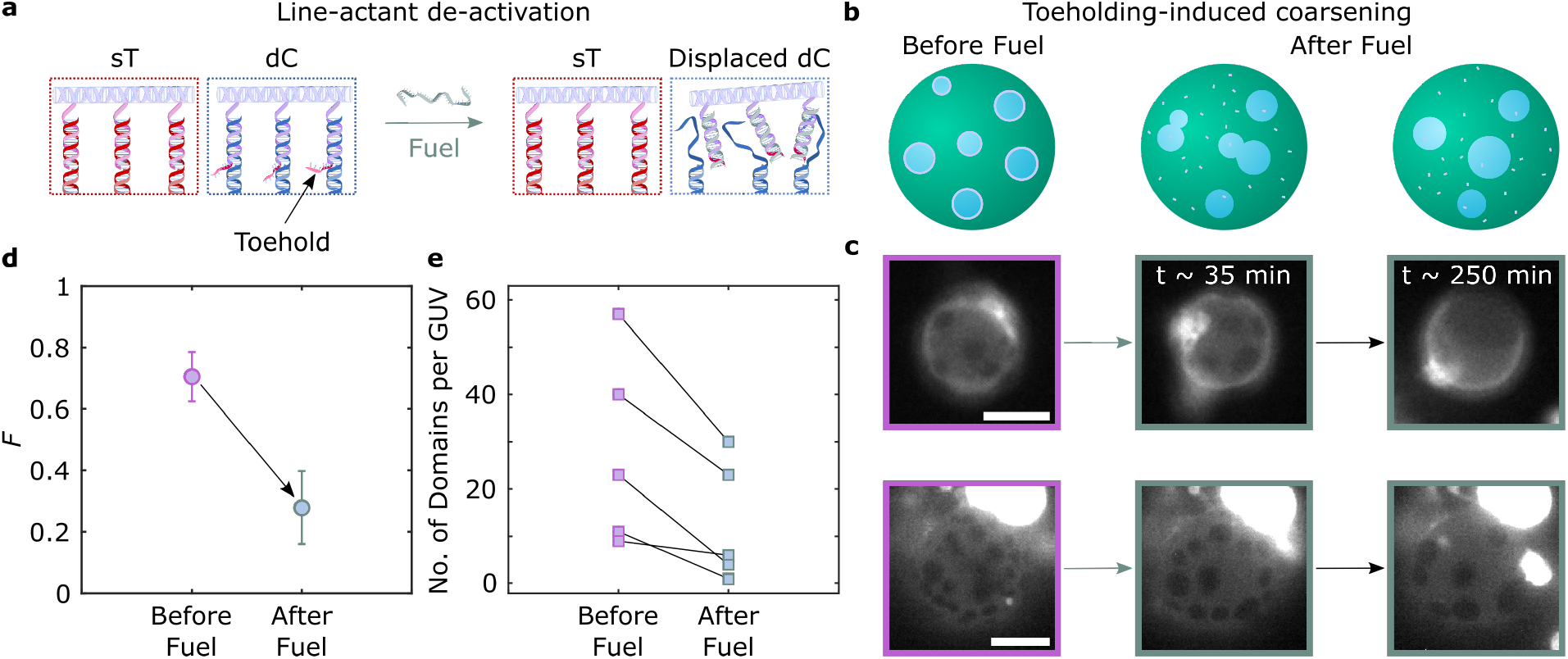
Toehold-mediated strand displacement enables dynamic control over membrane lateral organisation *via* DOLA reconfiguration. **a** Schematic depiction of anchor-points in DOLAs, highlighting the toehold domain linked to the dC-targeting overhangs. The addition of a Fuel strand catalyses a strand displace-ment reaction, detaching the DNA tile from the anchor-points and leading to DOLA de-activation. **b** DOLA re-configuration, and their subsequent desorption from line-interfaces, lead to domain coarsening. **c** Representative epi-fluorescence micrographs acquired before and after (∼ 35 and ∼ 250 minutes) the addition of Fuel, demonstrating domain coarsening triggered by DOLA de-activation. Fluorescent marker is Texas Red-DHPE, which partitions to the *L*_d_ phase. Scale bars = 10 *μ*m. **d** Influence of DOLA re-configuration on the fraction, *F*, of GUVs exhibiting more than two domains with respect to that of Janus-like vesicles before (*n* = 60 vesicles) and after (∼ 250 minutes, *n* = 74 vesicles) the addition of Fuel. **e** Number of domains per GUV in 6 representative vesicles, showing the effect of toeholding before and after (∼ 250 minutes) the addition of Fuel. Points connected by lines are relative to the same GUV.

As sketched in Fig. 5b (left), and shown in Fig. 5c (left) with fluorescence micrographs, GUVs decorated with the responsive 12× tiles form stable 2D emulsions when subjected to the heating-cooling cycle outlined above. Figures 5b–c (centre and right) show domain coarsening triggered by exposure to the Fuel strand, which translates into a clear decrease in the fraction *F* of non-Janus GUVs, shown in Fig. 5d. The series of snapshots in Fig. 5c shows that, while some GUVs acquire a Janus morphology after Fuel exposure (top), others retain a larger number of domains. Nonetheless, all observed GUVs experience a substantial degree of domain coarsening, as outlined in Fig. 5e where we track changes in the number of domains following Fuel addition in individual GUVs. As exemplified in Supplementary Fig. 15, domains often bulge after fuel addition, possibly due to a slight osmolarity mismatch coupled with differences in spontaneous curvature between the phases. Bulging may in turn be responsible for enhancing stability against coalescence of the domains after line-actant de-activation, preventing some GUVs from relaxing into the Janus configuration. ^74^

### Responsive DOLAs for domain fission in synthetic membranes

Reversible domain stabilisation with DOLAs can be exploited to synthetically replicate the action of biological membrane-remodelling machinery, tackling a critical bottleneck in synthetic-cell engineering. ^75^ For instance, phospholipase A_2_ has been postulated to exhibit enhanced catalytic activity at line-interfaces between co-existing *L*_d_–*L*_o_ phases, driving domain budding and fission.^76^ We propose our line-actants could form the basis of an analogous pathway coordinating fission and, therefore, three-dimensional membrane transformation.

We prepared GUVs in an initial configuration featuring DOLA-stabilised micro-domains, as outlined above. We then applied an hyperosmotic shock by increasing the osmolarity of the outside solution (*C*_O_) to ∼ 1.28 *× C*_I_, where *C*_I_ is the internal vesicle osmolarity, thus leading to an increase in GUV excess area.^77^ While in some cases hyperosmosis led to membrane internalisation (Supplementary Fig. 16), some domains responded by bulging-out, as depicted in Fig. 6a and experimentally observed in Fig. 6b. At this stage, some bulged-out domains could be “primed” for complete budding-off or fission, driven by minimisation of line-interface energy and opposed by membrane fluctuation entropy^78^ and, possibly, steric repulsion between DOLAs preventing line shrinkage. De-activation of the line-actants through Fuel strand addition is expected to increase line tension and negate any steric hinderance to interface shrinkage, tipping the marginally stable domains into full budding off and fission, (Fig. 6c, top). Fission events can be promptly recorded experimentally upon Fuel-strand addition, as shown in Supplementary Movie 6 and Fig. 6c (bottom). The sequence of epifluorescence images in the latter clearly shows the dark region corresponding to a budding *L*_o_-domain shrinking progressively, and ultimately disappearing. Vesicle fission is confirmed with brightfield micrographs, where arrows highlight both the “parent” and “offspring” vesicles regaining spherical morphologies. An analogous example is provided in Fig. 6d, and the associated Supplementary Movie 7, with overlaid confocal and bright-field micrographs.

**Figure 6:**
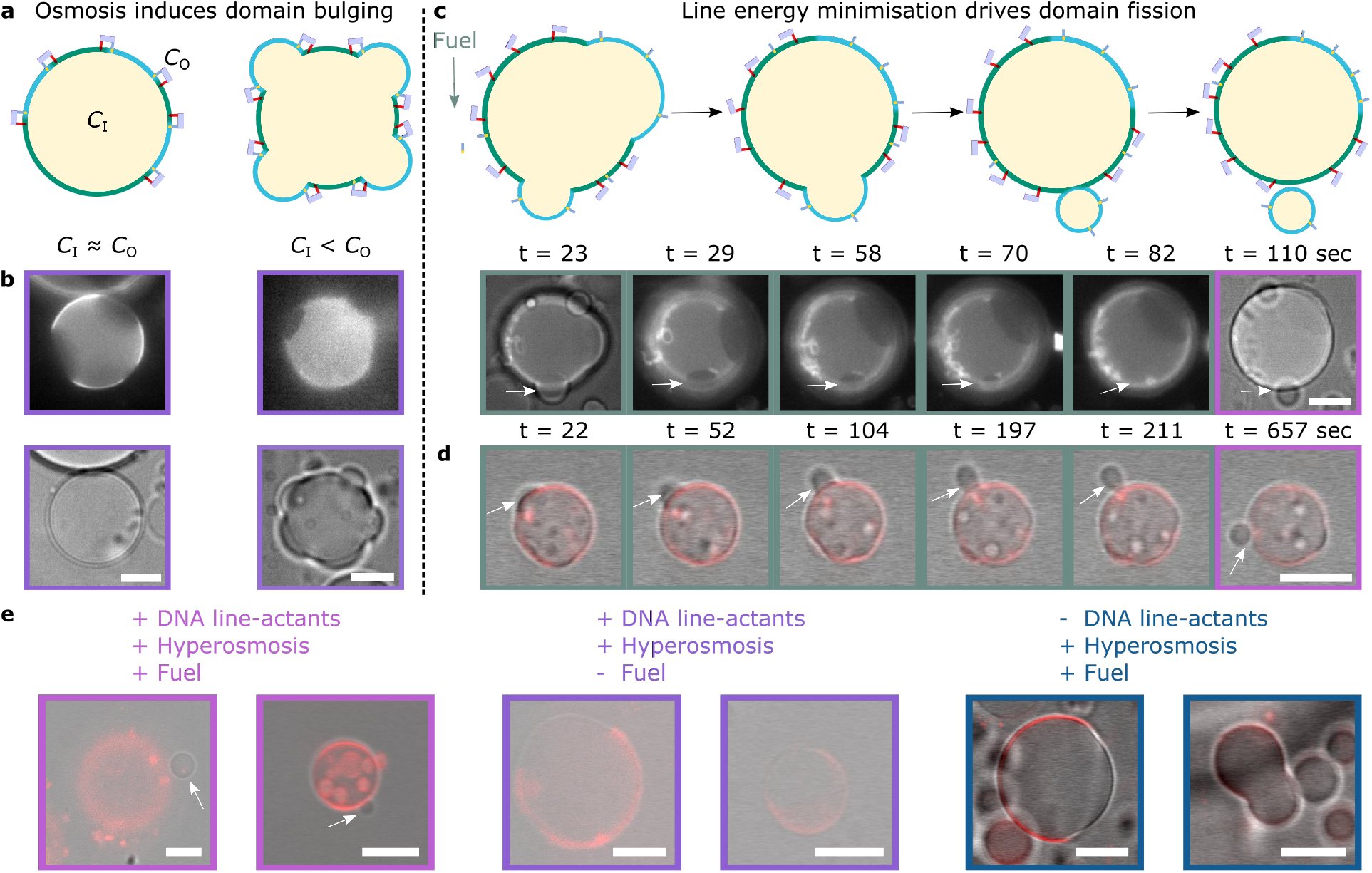
Biomimetic fission pathway mediated by DOLAs. The fission mechanism requires two sequential processes: domain budding and domain fission. **a** Schematic representation of the initial step, where bulging-out of DOLA-stabilised domains is caused by hyperosmotic shock, *i*.*e*. by increasing the outer concentration of solutes (*C*_*O*_) relative to the concentration inside the GUVs (*C*_*I*_). **b** Epifluorescence (top) and brightfield (bottom) micrographs of representative GUVs exposed to iso-osmolar (*C*_*I*_ ≈ *C*_*O*_) or hyper-osmolar (*C*_*I*_ *< C*_*O*_) environments, showing respectively spherical or bulging-out morphologies. **c** (Top) Schematic depiction of the second step, where Fuel addition and tile desorption from the interfaces induce budding off or fission of the protruding domains. (Bottom) Evolution of *L*_o_−domain fission in a representative liposome observed with brightfield and epifluorescence microscopy (see Supplementary Movie 6). Time-stamps refer to the time elapsed after initial acquisition. Fuel was added ∼ 2 minutes before acquisition began. **d** Evolution of *L*_o_ −domain fission in a representative GUV shown with overlaid confocal and bright field images (see Supplementary Movie 7). Time-stamps refer to the time elapsed after initial acquisition. Fuel was added ∼ 90 minutes before acquisition began. **e** Overlaid confocal and bright field micrographs of representative liposomes when exposed to: hyper-osmotic environments containing Fuel (Left) or lacking Fuel (Middle), or in the presence of Fuel but lacking DNA line-actants (Right). Fluorescent marker is Texas Red-DHPE, which partitions to the *L*_d_ phase. All scale bar = 10 *μ*m.

In further support of our proposed synthetic fission pathway, DOLA-functionalised liposomes with stabilised micro-domains were gently added to imaging wells containing hyperosmotic buffer baths with and without Fuel. In the former scenario, confocal microscopy revealed further examples of phase-separated GUVs found in proximity of a smaller single-phase vesicle (Fig. 6e, left, and Supplementary Fig. 17), similar to the confirmed fission occurrences shown in Fig. 6c and d. In turn, we did not observe such presumed fission events in vesicles immersed in baths lacking Fuel (Fig. 6e, middle), where GUVs appeared more Janus-like, also in line with our findings on sample heterogeneity summarised in Supplementary Fig. 16. Finally, DOLA-lacking GUVs, immersed in Fuel-supplemented hyper-osmotic buffer, simply adopted dumb-bell shapes thanks to their Janus-morphology (Fig. 6e, right), consistent with the expectation that the used *C*_O_*/C*_I_ is insufficient to drive fission of quasi-hemispherical domains.^79^

## Conclusions and Outlook

In summary, we presented DNA origami nanostructures – dubbed DOLAs – designed rationally to accumulate at the one-dimensional line-interface between lipid domains, imitating biological line-active membrane inclusions. The synthetic line-actants exploit the programmability and modularity of DNA origami, combined with the natural tendency of cholesterol and tocopherol “anchors” to enrich distinct lipid phases. Through experiments and numerical modelling, we demonstrated that the affinity for line-interfaces can be programmed by tuning the number of hydrophobic anchors, thus outlining a general design principle readily applicable to different origami designs, hydrophobic groups, or membrane compositions. We further demonstrated that our line-actants are able to stabilise small lipid domains against coarsening, giving rise to two-dimensional analogues of Pickering emulsions on the surface of giant liposomes, and achieving a previously unattainable degree of control over the lateral organisation of synthetic bilayers.

The nano-devices can be externally de-activated through a toehold-mediated strand displacement operation, triggering domain coalescence. Coarse-grained Monte-Carlo simulations replicate the experimental phenomenology, identifying origami-origami steric repulsion as the primary mechanism leading to domain stabilisation, and ultimately validating our line-actant engineering strategy. Finally, we combined externally-triggered line-actant de-activation with osmotic unbalance to implement a proof-of-concept synthetic fission pathway, whereby offspring liposomes programmably bud-off from parent vesicles, demonstrating the ability of the nano-devices to promote large-scale membrane re-structuring similar to biological nano-machines.^23,80,81^

The simplicity and modularity of the mech-anism underpinning line-action in our devices points at possible design variations that exploit different origami geometries, anchor chemistry, and polymerisation approaches to further program the number, size and morphology of lipid domains, marking a conceptual shift in synthetic-membrane engineering. Responsiveness to a wider range of physical and chemical stimuli can also be achieved by incorporating active elements such as DNAzymes,^82^ G-quadruplexes, ^83^ aptamers,^84^ and/or photoactuable moieties (e.g. azobenzene),^85^ thus broadening the design space and applicability of synthetic membrane-restructuring pathways. Thanks to their ability to program membrane re-structuring and the lateral distribution of membrane inclusions, the DOLAs constitute a valuable toolkit in synthetic-cell science that could underpin elusive and highly sough-after behaviours such as synthetic-cell division, vesicle-based trafficking and signal transduction pathways, ultimately unlocking disruptive applications to advance therapeutics and diagnostics.

Finally, DOLAs could be used to exert control over the local domain structure of living-cell membranes. One could thus envisage applying the origami line-actant toolkit as the basis of biophysical studies aimed at clarifying the role of micro-phase separation in cell signalling, by triggering the formation and disruption of “lipid rafts” at will. ^20^ If strong correlations between disease-related pathways and membrane-domain structure are established, the origami line-actants could carry potential therapeutic value by unlocking a previously unattainable strategy for conditioning cell behaviour.

## Acknowledgement

R.R.S. acknowledges support from the EPSRC CDT in Nanoscience and Nanotechnology (NanoDTC, Grant No. EP/L015978/1), the Mexican National Council for Science and Technology and the Cambridge Trust. L.D.M. acknowledges support from a Royal Society University Research Fellowship (UF160152, URF R 221009). R.R.S. and L.D.M. also acknowledge funding from the Royal Society Research Fellows Enhanced Research Expenses (RF/ERE/210029) and from the European Research Council (ERC) under the Horizon 2020 Research and Innovation Programme (ERC-STG No 851667 NANOCELL). The authors acknowledge computational resources provided by the Consortium des Équipements de Calcul Intensif (CÉCI), funded by the Fonds de la Recherche Scientifique de Belgique (F.R.S.-FNRS) under Grant No. 2.5020.11 and by the Walloon Region. A dataset in support of this work can be accessed free of charge at: https://doi.org/10.17863/CAM.94868

## Supporting Information Available

The Supporting Information is available at:

The following file is available free of charge including:

- Experimental Methods
- Supplementary Notes
- Supplementary Figures
- Supplementary Table with parameters for model and simulations
- DNA sequences of the nanostructures used throughout this work.

## TOC Graphic

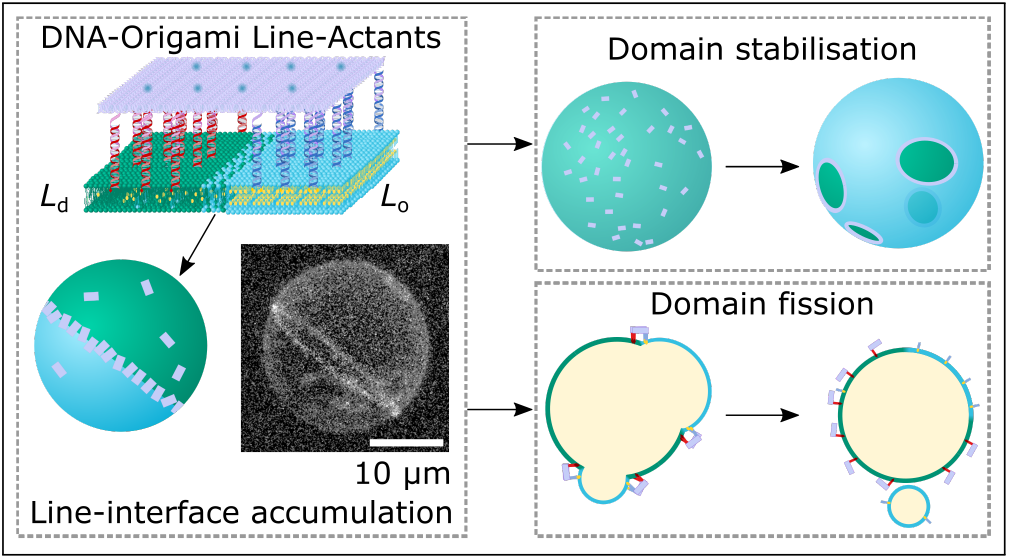

