## Supporting Information for "DNA-origami line-actants control domain organisation and fission in synthetic membranes"

### Methods

#### Electroformation of Phase-separating Giant Unilamellar Vesicles

Electroformed vesicles[1,2] were prepared in a 300 mM sucrose solution in MilliQ water as reported previously.[3] Briefly, lipid mixtures were composed of 1,2-dioleoyl-sn-glycero-3-phosphocholine (DOPC, chain melting temperature -17°C; Avanti Polar Lipids), 1,2-dipalmitoyl-sn-glycero-3-phosphocholine (DPPC, chain melting temperature 41°C; Avanti Polar Lipids), and cholestanol (Sigma-Aldrich) at 4:4:2 molar ratio. In addition, fluorescent Texas Red-DHPE (Invitrogen) was included in the lipid mixtures at 0.8% molar ratio, which preferentially enriches $L_\mathrm{d}$ domains. Subsequently, 45 $\mu$L of lipid mixtures (4 mg· mL$^{-1}$) were spread on the conducting side of clean indium tin oxide (ITO) slides heated above 60°C. The slides were placed in a dry silica dessicator for ∼1 hour and then assembled into an electroformation chamber using a ∼1 mm thick polydimethylsiloxane (PDMS) spacer enclosing approximately 400 $\mu$L of degassed sucrose buffer. Electroformation chambers were placed in a pre-heated oven at $T > 60$°C and connected to a frequency generator using clamps. The electroformation program consisted of a sinusoidal alternating current (AC), voltage amplitude of 2V, and frequency of 10 Hz for 2 hrs followed by 2 Hz for 1 hr. Finally, vesicles were retrieved and stored at room temperature in the dark to prevent photo-bleaching and photo-oxidation.

#### Design of line-active DNA origami nano-devices

DNA anchoring modules were designed using the NUPACK suite software,[4] and are based on previously reported nano-devices.[3,5] For the line-active plates, we used a modified version of the Rothemund Rectangular Origami.[6] Membrane attachment was achieved by functionalising the origami plates with the Picasso software,[7] which allowed to precisely engineer the spatial location of membrane-anchoring points. To that end, the sequence of specific staples was extended on their 3' end to include docking domains that would project outwards on a single face of the planar origami. These overhangs hybridised to sticky-end overhangs of dsDNA anchoring modules. On the opposite face of the origami, 5' end extensions were incorporated in eight DNA staples to include a domain capable of binding a fluorescent oligonucleotides for confocal imaging. In the case of line-partitioning measurements (Fig. 2 in the main text), 5'-end extensions were used to attach a strand with 3× binding sites for fluorescent oligonucleotides, in order to increase the signal-to-noise upon confocal acquisition.

#### Assembly of DNA Nanostructures

Lyophilised DNA oligonucleotides (Integrated DNA Technologies [IDT], Eurogentec, and Biomers), were reconstituted to a nominal concentration of 100 $\mu$M in Tris – Ethylenediaminetetraacetic acid (EDTA) buffer (1× TE: 10 mM Tris + 1 mM EDTA, pH 8.0).

DNA anchor-modules were subjected to a slow quenching temperature ramp on a TC-512 thermal cycler (95°C down to 4°C at a rate of –0.5°C min $^{-1}$) in a buffer containing 1× TE + 100 mM NaCl.

Origami plates self-assembled with a temperature quenching ramp in a total volume of 100 $\mu$L containing 7249-bp single-stranded circular viral template (M13mp18, tilibit nanosystems) as scaffold (10 nM) as well as core and modified stapling oligonucleotides, in the presence of a folding buffer composed of 1× TE + 12.5 mM $MgCl_2$. Staples capable of membrane-attachment were included at 100× excess, while core staples as well as those part of the fluorescent beacon were present at a 10× excess.

Thermal annealing was performed in a TC-512 thermal cycler. Samples were incubated at 80°C for 5 minutes to promote denaturing and suppress secondary structures. Subsequently, the temperature was decreased to 60°C, and later on to 40°C with a rate of $0.3125°C \cdot min^{-1}$. Finally, the temperature was brought down to 20°C.

#### Purification of DNA origami

Self-assembled rectangular DNA origami plates underwent 3 cycles of PEG-induced precipitation[8] to remove excess staples. To that end, DNA origami samples were brought to a volume of 250 $\mu$L with folding buffer (1× TE + 12.5 mM $MgCl_2$) and mixed 1:1 to a final volume of 500 $\mu$L with a solution containing 8000 Da polyethylene glycol (PEG8000, Sigma-Aldrich) at 15% (w/v) in 1× TE + 505 mM NaCl. After mixing *via* tube inversion, the solution was centrifuged (16,000g) at room temperature for 25 minutes. The supernatant was discarded, the pellet was reconstituted in 250 $\mu$L folding buffer, and the solution incubated at room temperature for at least 30 minutes before proceeding with the next precipitation cycle. After 3 cycles, the origami pellet was reconstituted to a final concentration of 10 nM and stored at room temperature in the dark to prevent photobleaching.

#### Vesicle Functionalisation with DNA line-actants

Pre-assembled anchoring modules were incubated overnight with vesicles (16.7 $\mu$L of DNA solution + 9.2 $\mu$L GUV solution, both prepared as indicated in "Electroformation of Phase-separating Giant Unilamellar Vesicles" and "Assembly of DNA Nanostructures", respectively, in 57.4 $\mu$L of buffer resulting in an iso-osmolar mixture. This functionalization procedure has been previously used in several works, and ensures optimal attachment of DNA constructs to GUVs[3,9–13]. Following the functionalisation of GUVs with anchor modules, the purified DNA origami tiles were heated up to $\sim$ 40°C for 10 minutes, followed by gentle vortexing to destabilise aggregates, and 8.35 $\mu$L were mixed with a second correction buffer (10 $\mu$L). The mixture was gently added to the GUVs decorated with anchoring modules, resulting in an iso-osmolar solution (1× TE + 99.2 mM NaCl + 0.8 mM $MgCl_2$ + 86.2 mM Glucose) that was left under rotation for at least 3 hours to allow for origami hybridisation to the anchor modules, and hence attachement of the tiles to the GUVs. In all cases, the membrane-bound anchor modules were in a nominal $\sim$ 1.5× excess of the origami binding capacity, to ensure that all avaialble overhangs on the origami are linked to the corresponding anchoring modules. Finally, the GUVs were placed in silicone wells for imaging, and allowed to sink for at least ∼15 minutes before imaging.

#### Agarose Gel Electrophoresis (AGE)

Agarose gels were prepared at 2% (weight) in Tris-Borate-EDTA (TBE, Sigma-Aldrich, 89 mM Tris-borate, 2 mM EDTA, pH 8.3) buffer. The mixture was dissolved *via* heating. Subsequently, SYBR Safe DNA gel stain (Invitrogen) was added at 0.1% (volume) mixed through gentle swirling, and was later casted to a thickness of approximately 5 mm. After setting for 1 hour, the gel was placed in an electrophoresis chamber and covered with TBE. 20 $\mu$L of annealed origami (Supplementary Fig. 1) or anchor-module samples (Supplementary Fig. 13) were loaded onto the gel alongside a DNA reference ladder (100 bp GeneRuler, Thermo Scientific). Control samples included origami staples and annealed scaffold. The electrophoresis chamber was placed on ice, and a potential of 75 V (3.75 V cm$^{-1}$) was applied for 90 minutes. Finally, the gel was imaged with a GelDoc-It system containing a UV lamp for illumination.

#### Fluorescence Microscopy of GUVs

Clean glass slides (15 min sonication cycles Hellmanax III 2%/isopropanol/MilliQ water) were passivated by covering with a solution containing bovine serum albumin (BSA) at 0.1% (w/v) and incubating in a pre-heated oven ($T > 60$°C) for an hour. Excess of BSA was removed with thorough rinsing with milli Q water. Silicone incubation chambers (Sigma-Aldrich) were stuck to passivated slides and sealed with DNAse-free tape to prevent evaporation.

Confocal imaging was performed at 1400 Hz, averaging over 10 frames, with a Leica TCS SP5 confocal microscope and an HC PL APO CORR CS 40× / 0.85 dry objective from

Leica. Alexa488 (excitation maximum - 495 nm; emission maximum - 520 nm) signal was acquired exciting with an Ar-ion laser (488 nm).

Epifluorescence microscopy was carried out at 10 frames per second using a home-built Nikon Eclipse Ti-E inverted microscope equipped with a 40× objective lens from Nikon (Plan APO $\lambda$, NA 0.95) and a Grasshopper3 GS3-U3-23S6M camera from Point Gray Research. Illumination in this set-up was provided by single-colour light emitting diodes (LEDs) through a filter set for Texas Red.

#### Atomic Force Microscopy

Atomic Force Microscopy (AFM) was used to confirm the correct assembly of DNA origami plates. After the temperature ramp, $10\,\mu$L of DNA origami samples at $\sim 1\,$nM (1× TE + 12.5 mM $MgCl_2$) were casted on top of a freshly cleaved mica sheet, which was previously fixed onto a microscope slide and cleaved with sticky-tape. The sample was incubated for 10 minutes before two cycles of washing. These consisted of adding $300\,\mu$L of ultrapure water on top of the mica sheet, followed by drying under a gentle nitrogen flow for 3 minutes. Micrographs were acquired using a MFP-3D Infinity AFM (Asylum Research) in Dry Tapping Mode. AFM silicon probes (BudgetSensors) bearing an aluminum reflex coating had a nominal frequency of $\sim 300\,$kHz and a stiffness of $40\,\mathrm{N} \cdot \mathrm{m}^{-1}$. Data processing was performed with Gwyddion[14] by levelling the data and applying mean plane subtraction and row alignment.

### Supplementary Note I: Membrane biophysics considerations for line-actant engineering

It is key to our line-actant design to compare the size of the DNA nanostructures with the finite thickness of the interface between two phases, and to ensure that same-type anchor-points are close together and separated from those with opposite partitioning tendency by a distance that exceeds the interface width. To that end, we estimated the thickness of the interface using the correlation length ($\xi$), derived as[15]

$$\xi \approx A \left| \frac{T - T_\mathrm{c}}{T_\mathrm{c}} \right|^{-\nu} \tag{S1}$$

where $\nu$ is a critical exponent, $A$ is an amplitude factor, $T$ is the temperature, and $T_\mathrm{c}$ is the critical temperature, defined as that where membranes with a suitable composition $\Phi_\mathrm{c}$ exhibit fluctuations with length scales that diverge and the line tension vanishes. The critical exponent has been shown, within experimental error, to be consistent with the 2-D Ising model prediction of $\nu = 1$,[15] while the amplitude factor ($A$) is given by $A = f^{(-)} \cdot \xi_0$ for $T < T_\mathrm{c}$, with $f^{(-)} \approx 0.17$ as a dimensionless factor, and $\xi_0 \approx 1\,\mathrm{nm}$ ( approximately the diameter of a lipid head group).[16]

In our implementation, for phase-separating membranes with a miscibility transition temperature ($T_\mathrm{m}$) close to the critical temperature, $T_\mathrm{m} \sim T_\mathrm{c}$ can be assumed. Indeed, we worked with vesicles prepared at 4:4:2 (DOPC/DPPC/Chol), whose $T_\mathrm{m}$ ($\sim 33\pm1$°C) has been shown to change only slightly with composition,[2] thus suggesting proximity to the critical point ($\Phi_\mathrm{c}, T_\mathrm{c}$). It is therefore possible to obtain a sufficiently good approximation of $\xi$ at room temperature ($T \sim 25^\circ$ C) with Eq. S1 using $T_\mathrm{m} \approx T_\mathrm{c}$, leading to an estimated width of the interface of $\sim 8\,\mathrm{nm}$.

### Supplementary Note II: Data acquisition and Data analysis

#### Data Acquisition

DNA-functionalised GUVs were placed in silicone incubation chambers and allowed to sediment on the bottom for at least 10 minutes prior to imaging. The chambers were sealed with sticky-tape to prevent evaporation.

##### Confocal Microscopy

Confocal micrographs were processed to assess the line-accumulation tendencies of our DNA nano-devices. To that end, we acquired z-stacks of fields of view $64.84\,\mu m \times 64.84\,\mu m$ with slice thicknesses of $0.5\,\mu m$. The size of the field of view is the largest our instrument can image at the fastest scanning rate of 1400 Hz. Screening and selection of imaged vesicles was underpinned by two criteria:

1. Vesicles were smaller than the size of the field of view and bigger enough such that Brownian motion was negligible within the acquisition timescales of a few minutes per GUV.

2. The orientation of the vesicle was such that a sharp interface separating the $L_\mathrm{d}$ and $L_\mathrm{o}$ domains was visible at the equatorial cross-section of the GUV, thus enabling quantitative processing.

In view of the size cut-off introduced by the first criterion, resulting in an experimental relevant range of radii of $5 < R < 25\,\mu m$, our model has accounted for the influence of GUV radius on the physics of line-accumulation.

##### Epifluorescence Microscopy

Epifluorescence micrographs were acquired to measure the ability of DNA line-actants to stabilise domains against coalescence. Passivated slides with GUV-containing imaging chambers were placed on top of a copper plate connected to a Peltier element and fixed with aluminum tape. The chambers were then heated to $T = 37°C$, allowed to equilibrate for $\sim 5$ minutes, and cooled down to $T = 25°C$. The chambers were subsequently placed on the microscope stage, and the vesicles were allowed to equilibrate for $\sim 3$ hours. Acquisition was done using a custom-built script that enabled the automated manipulation of the instrument in terms of time and illumination at previously hand-picked positions.

#### Data Analysis

##### Quantitation of line-accumulation tendencies across variants of line-active DNA nano-devices

Equatorial micrographs of GUVs were manually selected from confocal z-stacks prior to analysis. Image processing is underpinned by a MATLAB segmentation pipeline that allows to identify membrane radial intensities and extract data points around the boundaries between lipid domains.

The custom-built user-assisted software loads equatorial micrographs and uses a Gaussian blurring routine for noise removal and to facilitate segmentation. Aided by a Graphical User Interface, three positions along the membrane circumference are manually chosen: i and ii) are rough locations of the two interfaces and iii) is a position along the $L_\mathrm{d}$ domain. Estimated values are subsequently used to enclose the membrane within a ring, allowing the software to extract radial profiles along the membrane contour in steps of $\pi/900$. For each profile, the position of the membrane is estimated using a Gaussian fit. Estimated positions are then processed with a circle-fitting routine. Specifically, a subset of points extracted from radial profiles (those within $\pm 5\%$ of the estimated average radius of the vesicle) are fitted to a circle, which we used as a final estimate for the membrane positions. These enabled us to extract from the unprocessed raw images the average fluorescence intensity of the membrane *via* numerical integration around the position of the bilayer ($\pm 2$ pixels).

We subsequently extracted segments from the radial fluorescence intensity profile corresponding to the line-interfaces *via* the user-provided rough locations of the two boundaries. The traces were extracted around a defined width of $\pm 5\,\mu m$ from the estimated location of the interface, which we then fitted to an empirical model to better approximate the location of the line-interface. To that end, we model the fluorescence profile of the $L_\mathrm{d}$-$L_\mathrm{o}$ interface as the sum of a step function with a rectangular function, both centred at the location of the boundary. Owing to diffraction-limited blurring, the sum of these functions is convolved with a Gaussian.

We thus fit our fluorescence intensity data to the sum of a Gaussian and a sigmoidal function as

$$I(x) = A\mathrm{exp}\left[-\left(\frac{x-B}{C}\right)^2\right] - D\mathrm{tanh}\left[\frac{x-B}{C}\right] + D + E \tag{S2}$$

where $A$ and $D$ are the amplitude of the Gaussian and sigmoidal, respectively, and C is the width. From the fit, we estimate the location at which the interface is centred, namely $B$. Finally, vector re-sampling is applied to construct an interfacial intensity trace around the location of the interface ($x = 0$) around a width of $\pm 4\,\mu m$ at every $0.05\,\mu m$. Line-interface traces were then averaged and fitted to Eq. S2, as collated in Fig. 2 (main text) and Supplementary Fig. S8.

From the fitted parameters $A, C, D$, and $E$, we compute the line partitioning coefficient $K_\mathrm{p,int}$. First, the bulk $L_\mathrm{d}$ fluorescence intensities were estimated as $I_{L_\mathrm{d}} = D+E$ and $I_{L_\mathrm{o}} = E$. Then, the average fluorescence intensity of the line-interface ($I_\mathrm{int}$) was approximated with

$$I_{\text{int}} = \frac{(A+E)C\sqrt{\pi}}{\delta} \tag{S3}$$

In other words, we calculated the area under the Gaussian (i.e. the fluorescence intensity across the whole finite thickness of boundary) and divided it by the nominal length along which the origami can accumulate (i.e. $\delta = \xi + \ell$, where $\ell$ is the spacing between dC and sT anchoring regions in the origami).

We then define a mean number density $\rho = N/A_{\text{GUV}}$ as the total number of origami in a vesicle with surface area $A_{\text{GUV}}$. Similarly, it is possible to define mean number densities at the line-interface ($\rho_{\text{int}} = N_{\text{int}}/A_{\text{int}}$) and in $L_{\text{o(d)}}$ phases ($\rho_{L_{\text{o(d)}}} = N_{L_{\text{o(d)}}}/A_{L_{\text{o(d)}}}$).

Thus, line-interface partition coefficients can be obtained as

$$K_{\text{p,int}} = \frac{\rho_{\text{int}}}{\rho} \tag{S4}$$

Assuming that the measured fluorescence intensities are proportional to the membrane densities, such that $\rho_{\text{int}} = \alpha I_{\text{int}}$ and $\rho_{L_{\text{o(d)}}} = \alpha I_{L_{\text{o(d)}}}$, we can write

$$\rho = \alpha \frac{I_{\text{int}}A_{\text{int}} + I_{L_{\text{o}}}A_{L_{\text{o}}} + I_{L_{\text{d}}}A_{L_{\text{d}}}}{A_{\text{GUV}}} \tag{S5}$$

which applied to Eq. S4 leads to

$$K_{\text{p,int}} = \frac{I_{\text{int}}A_{\text{GUV}}}{I_{\text{int}}A_{\text{int}} + I_{L_{\text{o}}}A_{L_{\text{o}}} + I_{L_{\text{d}}}A_{L_{\text{d}}}} \tag{S6}$$

We used Eq. S6 to compute the experimental line-partitioning coefficients, collated in Supplementary Fig. S5. To that end, we assumed the vesicles (with radius $R$) were perfectly Janus with two hemispherical domains, thus having $A_{\text{GUV}} = 4\pi R^2$, $A_{L_\text{o}} = A_{L_\text{d}} = 2\pi R^2$. In turn, we estimate the area of the interface using $A_{\text{int}} = 2\pi R\delta$, i.e. the length of the line-interface ($L = 2\pi R$) multiplied by $\delta = \xi + \ell$ (the width available for tile displacements in the direction orthogonal to the line-interface). Recalling that from the fit with Eq. S2 we estimate $I_{\text{int}} = \frac{(A+E)C\sqrt{\pi}}{\delta}$, $I_{L_\text{d}} = D + E$, and $I_{L_\text{o}} = E$, Eq. S6 becomes:

$$K_{\text{p,int}} = \frac{2R\frac{(A+E)C\sqrt{\pi}}{\delta}}{(A+E)C\sqrt{\pi} + R(D+2E)}. \tag{S7}$$

To compute the values of $K_{\text{p,int}}$ shown in Fig. 2**c**, we chose $\xi = 8\,\text{nm}$ (see Supplementary Eq. S1) and $\ell = 30\,\text{nm}$, for 6× and 12× tiles, or $\ell = 25\,\text{nm}$ for 24× tiles. While the latter value is defined by the origami design, substantial uncertainty may affect the estimate of the interface thickness $\xi$. To determine how this uncertainty propagates to the estimation of $K_{\text{p,int}}$, in Supplementary Fig. 8 we show the results obtained using the lower and upper physical bonds of $\xi$, with the lower bound being $\xi = 0$ ($\delta = \ell$, negligible interface thickness) and the upper bound being $\xi = \ell$ ($\delta = 2\ell$, determined by noting that $\xi > \ell$ would prevent line accumulation). $K_{\text{p,int}}$ values are, for all choices of $\xi$, within a factor two of each other and preserve the same trends with respect to tile design.

**Quantification of lipid domains stabilisation**

To assess the line-action of DNA nanostructures, DNA-decorated vesicles were imaged after temperature quenching using epifluorescence microscopy. Micrographs of several fields of view were acquired for each sample, and individual micrographs were analysed using a custom-built MATLAB script for quantitative assesment.

The script, which operates in a user-assisted fashion, loads a frame selected at random from a set containing images for all conditions (Fig. 3b in the main text), thus blinding the data and minimising human bias. The Graphical User Interface then asks the user to input values for:

- The number of vesicles that are de-mixed ($N_{\mathrm{DM}}$).
- The number of vesicles that are homogenously mixed ($N_{\mathrm{M}}$).
- The number of vesicles with Janus morphology (i.e. with only two macroscopic domains) ($N_{\mathrm{J}}$).
- The number of vesicles with two-dimensional Pickering emulsions (i.e. with more than two macroscopic domains) ($N_{\mathrm{PE}}$).

Once all frames have been allocated values for each of these conditions, the script computes the fraction of vesicles featuring more than two stable domains ($F$) with

$$F = \frac{N_{\mathrm{PE}}}{N_{\mathrm{PE}} + N_{\mathrm{J}} + N_{\mathrm{M}}}, \tag{S8}$$

### Supplementary Note III: Theoretical estimation of line-partitioning

Throughout this section, we derive a numerical model to predict line-partitioning of our DNA-origami line-actants. Thus, by comparing predictions from theory using experimental parameters (see Table S6) with the experimental observations (Figure 2 of the main text), we infer the surface coverage of tiles on GUVs ($\sigma$), which we subsequently use to compute the fraction of tiles at the line-interface ($\phi$).

We consider $N_{\rm int}$ tiles covering a line-interface of length $L$. The partition function of the system, $Z_{\rm int}$, accounts for the free-energy contribution given anchor–bilayer interactions ($\Delta G_{\rm p,int}$) as well as a term ($z_{\rm int,conf}$) accounting for the configurational space available to each tile. Thus,

$$Z_{\rm int} \quad = \quad \frac{\left[q_1 \exp(-\beta \Delta G_{\rm p,int}) z_{\rm int,conf}\right]^{N_{\rm int}}}{N_{\rm int}!} \; . \tag{S9}$$

Equation S9 assumes that lipid–lipid interactions are not altered by the presence of anchors. In addition, $q_1$ lumps all single–tile contributions with the exclusion of anchors–bilayer interactions. We write the configurational free–energy *per* tile as follows

$$z_{\rm int,conf} = \Delta\theta \left[\delta(L - N_{\rm int} L_1)\right] \; , \tag{S10}$$

In the previous expression, $\Delta\theta$ refers to the angle describing the range of possible tile orientations with respect to the line-interface, $\delta$ is the distance range available for possible tile displacements orthogonal to the line, and $L_1$ is the width (short side) of the tile. $\delta$ is estimated using the distance between the two sets of anchor-points ($\ell$) augmented by the finite width of the line-interface ($\xi$, see Fig. 1 in the main text and Supplementary Note I). We estimated $\Delta\theta$ by placing the centre of the tile on top of a straight line separating the two lipid phases, allowing us to consider all possible tile orientations with which all anchor-points are hosted on their favorable phase. Note that it is possible for a tile at the line-interface to expose some anchors to their unfavourable phase by paying an accessible free–energy cost (increasing the value of $\Delta G_{\rm p,int}$). Therefore, in order to estimate an order of magnitude of systematic errors associated with the model, below we also consider tiles which can fully rotate while being at the line (i.e., $\Delta\theta = 2\pi$). Note also that in Eq. S10 we factorize the translational (parallel and orthogonal to the line-interface) and rotational contributions while, in reality, rotational and translational degrees of freedoms are coupled (for instance, $\Delta\theta$ tends towards zero when two tiles enter into contact).

We then assess the equilibrium conditions between tiles in the bulk phases (modeled using two hemispheres with surface area $2\pi R^2$) and tiles at the line. We consider the dilute case in which, in the bulk phases, tile–tile interactions are negligible (note that Eq. S10 accounts for excluded volume interactions between tiles at the line-interface). Importantly, we do not consider the possibility for the tiles to increase the length $L$ of the line, which we fix to $L = 2\pi R$. We argue that this is a reasonable assumption at low $T$, as supported by the simulations in Fig. 4 (main text). The chemical potentials in the three different locations ($L_\mathrm{d}$, $L_\mathrm{o}$, and $L_\mathrm{d}$–$L_\mathrm{o}$ interface) are the same and equal to

$$\begin{aligned} \beta\mu &= \log \frac{N_\mathrm{int}}{(L - N_\mathrm{int}L_1)\delta q_1 \Delta\theta} + \frac{N_\mathrm{int}L_1}{L - N_\mathrm{int}L_1} + \beta\Delta G_{\mathrm{p,int}} \\ &= \log \frac{N_{L_\mathrm{o}}}{(2\pi R)^2 q_1} + \beta\Delta G_{\mathrm{p},L_\mathrm{o}} \\ &= \log \frac{N_{L_\mathrm{d}}}{(2\pi R)^2 q_1} + \beta\Delta G_{\mathrm{p},L_\mathrm{d}} \end{aligned} \tag{S11}$$

where $N_{L_\mathrm{o}(L_\mathrm{d})}$ and $\Delta G_{\mathrm{p},L_\mathrm{o}(L_\mathrm{d})}$ are, respectively, the number of tiles and the anchors-bilayer free energy contributions *per* tile in the $L_\mathrm{o}(L_\mathrm{d})$ phase. Equations S11 allow calculating $N_\mathrm{int}$. The problem can be simplified by assuming that the tiles at the boundary lines do not deplete the number of tiles in the bulk phases, $N_\mathrm{int} \ll N_{L_\mathrm{o}}, N_{L_\mathrm{d}}$, and therefore do not change the chemical potential of the bulk phases as compared to the $N_\mathrm{int} = 0$ case. Under these assumptions, using $N = N_{\mathrm{L_o}} + N_{\mathrm{L_d}}$ in the second equality of Equations S11 with $\rho = N/(4\pi R^2)$, we simplify the chemical potential as follows

$$\beta\mu = \log \frac{\rho/(q_1\pi)}{\exp(-\beta\Delta G_{\mathrm{p},L_\mathrm{o}}) + \exp(-\beta\Delta G_{\mathrm{p},L_\mathrm{d}})} \tag{S12}$$

The fraction of line-interface covered by tiles, assuming that tiles remain orthogonal to the line-interface, is

$$\phi = N_{\mathrm{int}} \frac{L_1}{L} \tag{S13}$$

which we then obtained using Eq. S12 in the first of the equalities in S11, which leads to

$$\frac{\phi}{1-\phi} \exp\left[\frac{\phi}{1-\phi}\right] = \frac{2L_1\delta\Delta\theta}{2\pi} \frac{\exp(-\beta\Delta G_{\mathrm{p,int}})}{\exp(-\beta\Delta G_{\mathrm{p},L_\mathrm{o}}) + \exp(-\beta\Delta G_{\mathrm{p},L_\mathrm{d}})}\rho \equiv \Lambda\rho \tag{S14}$$

from which $\phi$ can be calculated numerically. Note that, in a Langmuir–like approach that would not consider the second term in RHS of first equality in S11, the previous equation would result in

$$\phi = \frac{\Lambda\rho}{1+\Lambda\rho} \,. \tag{S15}$$

Given $\phi$, the theoretical prediction of the line–partitioning coefficient is calculated with

$$K = \frac{\rho_{\mathrm{int}}}{\rho} = \frac{\phi}{\rho L_1 \delta} \,. \tag{S16}$$

We now estimate the anchors–bilayer free-energy contributions $\Delta G_{\mathrm{p},L_\mathrm{o}}$, $\Delta G_{\mathrm{p},L_\mathrm{d}}$, and $\Delta G_{\mathrm{p},L_\mathrm{o}}$. The experimental $K_{\mathrm{p,int}}$ can be readily compared to the increase in surface density required to saturate line-interfaces. As done previously in our work on regulating the partitioning of DNA nano-devices in lipid domains,[3] we define a free-energy of partitioning as the shift associated with moving a single DNA construct from $L_\mathrm{d}$ to either the $L_\mathrm{d}$–$L_\mathrm{o}$ interface ($\Delta G_{\mathrm{p,int}}$) or $L_\mathrm{o}$ ($\Delta G_{\mathrm{p},L_\mathrm{o}}$). In particular, this choice sets $\Delta G_{\mathrm{p},L_\mathrm{d}}$ to $\Delta G_{\mathrm{p},L_\mathrm{d}} = 0$.

To achieve line-accumulation, dC anchor-points need to be translocated to $L_{\mathrm{o}}$ while maintaining sT modules in $L_{\mathrm{d}}$ so to induce the correct orientation of the line-actants. Thus, we calculate the free energy of line partitioning with $\Delta G_{\mathrm{p,int}} = \sum_i \Delta G^{i\mathrm{dC}}_{\mathrm{p},L_{\mathrm{o}}}$, where the index $i$ runs over all the dC anchors in the origami. In the case of enriching $L_{\mathrm{o}}$ domains, the free energy of partitioning is additive in the contributions of all ($j$) hydrophobic anchors featured in the nanostructure given by $\Delta G_{\mathrm{p},L_{\mathrm{o}}} = \sum_j \Delta G^{j}_{\mathrm{p},L_{\mathrm{o}}}$. Thanks to the work of Jungmann and co-workers,[17] one can also account for the efficiency of staple addressability in the origami by weighting the free-energy contribution of a given anchor-point (see Table S7) with the probability ($p$) of incorporation as

$$\Delta \tilde{G}_{\mathrm{p,int}} = \sum_i p_i \Delta G^{i\mathrm{dC}}_{\mathrm{p},L_{\mathrm{o}}} \tag{S17}$$

To account for possible systematic errors (as done for the rotational configuration contributions to the free–energy), also in view of the fact that we expect the line-interface to be enriched by tiles featuring more anchors, if not all, than the averaged values, in the following we consider both $\Delta G$ and $\Delta \tilde{G}$. All parameters employed in the theoretical model are summarised in Table S6. Note that the average incorporation probabilities for dC and sT anchoring staples are similar across the three tile designs adopted (see Table S8), implying that incomplete staple incorporation is unlikely to produce a substantial unbalance in the number of available binding sites available for the two types of anchoring modules.

Figure S5 (a,c, and e) report the theoretical predictions of the line-partitioning coefficient ($K_{\mathrm{p,int}}$) as a function of the surface coverage ($\sigma = \rho L_1 L_2$). By intersecting these plots with the experimental values $K^{n\times}_{\mathrm{p,int}}$ (including an uncertainty on the latter quantified with the standard deviation, $\delta K_{\mathrm{p,int}}$, see Table S6), we predict the following surface coverages, also summarised in Fig. S6: $\sigma^{6\times}_{\mathrm{theory}} < 0.032$ , $0.015 < \sigma^{12\times}_{\mathrm{theory}} < 0.093$, and $0.029 < \sigma^{24\times}_{\mathrm{theory}} < 0.144$. These values are consistent with the approximated coverage ($\sigma_{\mathrm{exp}} \sim 0.05$) using the stoichiometric parameters employed in our experimental implementation (see Methods in the main text). The increase of the predicted $\sigma$ in the 12× and 24× designs can be explained by a higher affinity of the tiles for the lipid membranes. This observation is consistent with the absolute values of the fluorescent signals reported in Fig. S8.

Using the inferred values of $\rho$, $\rho = \sigma/(L_1 L_2)$, we can then calculate the fraction of occupied line-interface ($\phi$) using Eq. S14 (see Fig. S5, panels b,d, and f). In particular, we find: $\phi^{6\times} < 0.537$, $0.508 < \phi^{12\times} < 0.84$, and $0.88 < \phi^{24\times} < 0.935$. These results show how the experimental values of $K_{\mathrm{p,int}}$ are indicative of the regulatory influence of anchor number in line-partitioning, demonstrating an increase in line-accumulation as a function of the number of anchors. Finally, our observations also prove that line–partitioning is mainly driven by the anchor–bilayer affinity, supporting the biophysical model proposed in this contribution.

Šulc and co-workers have recently demonstrated that 2D origami tiles spontaneously bend in the direction orthogonal to that of the double helices, as a result of steric repulsion between overhangs present only on one face.[18] Bending of our membrane-confined tiles would result in a lower effective width $L_1$, and thus impact the estimation of $z_{\mathrm{int,conf}}$.
Because of the even distribution of anchoring points on their surface, 24× and 12× tiles would need to break connections with the membrane in order to bend, or force a curvature on the lipid bilayer itself. These occurrences are however energetically prohibitive, given that the typical free energies associated with tile bending ($\sim 5k_{\mathrm{B}}T$[18]) are small compared with the free-energies penalties associated to i) de-hybridising the tile overhangs from the anchor modules tethered to the lipid bilayer ($\sim 40k_{\mathrm{B}}T$ per overhang, as computed with NUPACK[4]), ii) pulling out one of the hydrophobic anchors from the bilayer ($\sim$ 20 to $30k_{\mathrm{B}}T$ per anchor[19,20]), and iii) bending the membrane ($\sim 20k_{\mathrm{B}}T$ for liquid-disordered lipid membranes[21]). Due to the anchoring points being localised close to the long axis of the tile, 6× designs could potentially retain a bent configuration while anchored to the membrane. It should be however noted that Šulc and co-workers observed spontaneous curvature on

origami featuring a full array of 169 overhangs. We would thus speculate that the effect is likely negligible for the 6× design, which only has 12 overhangs.

We thus conclude that none of the tested tile designs are likely to acquire curved configurations while on the membranes, justifying the interpretation of $L_1$ as the nominal tile width.

### Supplementary Note IV: Implementation of Ising-type coarse-grained Monte Carlo simulations

#### The model

*Lipid bilayer.* Phase separation in lipid bilayers has been traditionally studied using the Ising model.[22–25] In particular, the thermodynamic quantities of the system and the line morphology close to the critical point are described by the Ising universality class.[15] A bilayer is modeled with a 2-dimensional lattice in which each site carries a fluctuating spin ($s_i$, $s_i = \pm 1$), representing the phase found in a unit cell ($L_\mathrm{o}$ or $L_\mathrm{d}$). Neighboring spins ($s_i$ and $s_j$) contribute to the energy of the system through a $-Js_is_j$ term, where $J$ is an energetic parameter favouring the alignment of the spins and, therefore, hampering the formation of boundaries between different phases. The total configurational energy then reads as

$$H_{ss} = -\frac{J}{2}\sum_i \sum_{j\in\nu(i)} s_i s_j \quad , \tag{S18}$$

where $\nu(i)$ is the list of neighboring sites of $i$. We use a triangular lattice, in which each site interacts with six neighbors, and employ periodic boundary conditions. The non-dimensional parameter controlling the phase behavior of the system is given by $x = J/(k_B T)$. In particular, phase separation is induced when decreasing the temperature below a critical value ($T_c$) corresponding to $x_c = \frac{\log 3}{4}$ ($T_c = \frac{J}{k_B x_c}$). The spin update implemented by the simulation algorithm (see below) conserves the total magnetization of the system defined as $M = \sum_i s_i$. The dynamics preserving $M$ are consistent with the conservation of the number of lipid molecules of a given type.

*Line-actants.* We now describe the tiles' model and their implementation to the Ising model introduced above (see Fig. 4a of the main text). $N_T$ is the number of tiles in the system. A tile covering a fraction of the bilayer is modeled by flagging a certain number of lattice sites as being covered by the specific tile. Accordingly, the position and orientation of the tiles are discretised variables. Following the design shown in Fig. 1b of the main text, each tile is approximated by 17 rows made of 15 lattice sites. As stated above, the rows are constrained to lay along one of the 6 directions of the lattice. A certain number of lattice sites covered by a tile will also host an anchor (shown in blue and red hexagons in Fig. 4a of the main text). The position of the anchors is fixed in the tile's reference frame. In particular, if the position of a tile is updated, the anchors will move accordingly.

We define by $i^{(\mathrm{sT})}_{k,\alpha}$ and $i^{(\mathrm{dC})}_{k,\alpha}$ ($k = 1, \cdots, N_A$, $N_A = 6$, 12, or 24) the lattice hosting the $k$–th anchor of tile $\alpha$ of type sT and dC, respectively. The interaction of a spin with neighboring spins (Eq. S18) is not altered by the presence of a tile (in particular a lattice site can be occupied both by a spin and an anchor). An anchor hosted by site $i$ interacts with $s_i$ and all the spins found on a neighbor of $i$. In particular, an anchor of type sT and dC will contribute to the configurational energy with $-J_{\mathrm{sT}}\tilde{s}(i^{(\mathrm{sT})}_{k,\alpha})$ and $-J_{\mathrm{dC}}\tilde{s}(i^{(\mathrm{dC})}_{k,\alpha})$, respectively, where we defined $\tilde{s}(i) = s_i + \sum_{j\in\nu(i)} s_j$. By taking $J_{\mathrm{sT}} > 0$ ($J_{\mathrm{dC}} < 0$) we design sT (dC) anchors that interact favorably with the $s = 1$ ($s = -1$) phase. Finally, the total contribution of the tiles to the configurational free energy reads as follows

$$H_t = -\sum_{\alpha=1}^{N_T}\sum_{k=1}^{N_A}\Big(J_{\mathrm{sT}}\tilde{s}(i^{(\mathrm{sT})}_{k,\alpha}) + J_{\mathrm{dC}}\tilde{s}(i^{(\mathrm{dC})}_{k,\alpha})\Big) + \sum_{\alpha<\beta} V^{(\mathrm{excl})}_{\alpha,\beta} \tag{S19}$$

where $V^{(\mathrm{excl})}_{\alpha,\beta}$ is an excluded-volume interaction preventing a lattice site to be occupied by more than one tile. Finally, the total configurational energy ($H$) of the model is given by

$$H = H_{ss} + H_t \quad . \tag{S20}$$

#### Estimation of the model's parameters

The line tension of spin systems on the triangular lattice is calculated as in Ref.[26] based on the method of Zandvliet to calculate boundary free-energies.[27] The free-energy cost *per* spin along one of the principal directions, $\tau^*$, reads as follows (see Eq. 9 in Ref.[26] with $A = 1$ and $B = X = Y = \exp(-2x)$)

$$\beta\tau^* = 4x + \log\frac{1 - e^{-4x}}{2} \tag{S21}$$

Note that $\tau^* = 0$ if $x = x_c$. The previous equation is similar to an exact result calculated by Onsager (see Eq. 122a in Ref.[28]) for the square lattice. By inverting the previous equation we obtain

$$x = \frac{1}{4}\log\left(1 + 2e^{\beta\tau^*}\right) \tag{S22}$$

We estimate $\tau^*$ as $\tau^* = \tau \cdot \delta$, where $\tau$ is an experimental estimation of the line tension ($\tau \approx 1 \cdot \mathrm{pN}$[15]) and $\delta$ ($\delta = 5.77\,\mathrm{nm}$) corresponds to the lattice spacing of the tile's geometry (see Fig. 1b, main text). Using these values in Eq. S22 provides $J = x\,k_B T \approx 0.55\,k_B T$.

The non-dimensional parameters controlling the interactions between the tiles and the lipid bilayer are $x_\mathrm{a} = \frac{J_\mathrm{a}}{k_B T}$ and $x_\mathrm{b} = \frac{J_\mathrm{b}}{k_B T}$. These parameters are calculated from the experimental values of $\Delta G_{L_\mathrm{o}}$. In particular, given that each anchor interacts with 7 spins, we have

$$14x = \beta|\Delta G_{L_\mathrm{o}}| \tag{S23}$$

from which, using the experimental values of $\Delta G_{L_o}$, we derive

$$J_{\mathrm{sT}} = 0.136\, k_B T \qquad\qquad J_{\mathrm{dC}} = -0.057\, k_B T\,. \tag{S24}$$

The previous equations neglect spin fluctuations (i.e. the fact that not all contacts could be favorable) and is strictly valid only in the limit in which temperatures tend to zero.

#### The simulation strategy

Each simulation cycle includes $N_a$ attempts to implement one of the following MC moves ($N_a = N_l + 2N_T$, where $N_l$ is the number of lattice sites and $N_T$ the number of tiles):
1–Spin update (executed with probability $p = 0.9$). Spins are updated using Kawasaki dynamics.[29] This move proposes to exchange a pair of neighboring spins and accepts the new configuration according to a Metropolis test[29] with the change in configurational energy calculated using $H_{ss}$.
2–Tiles update (performed with probability $p = 0.1$). With equal probability, we either try to move a randomly selected tile along one of the six lattice orientations by $\delta$, or we reorient the tile by $\pm 60°$. The proposed configuration is accepted according to a Metropolis test with the change in configurational energy calculated using $H_t$.

### Supplementary Figures

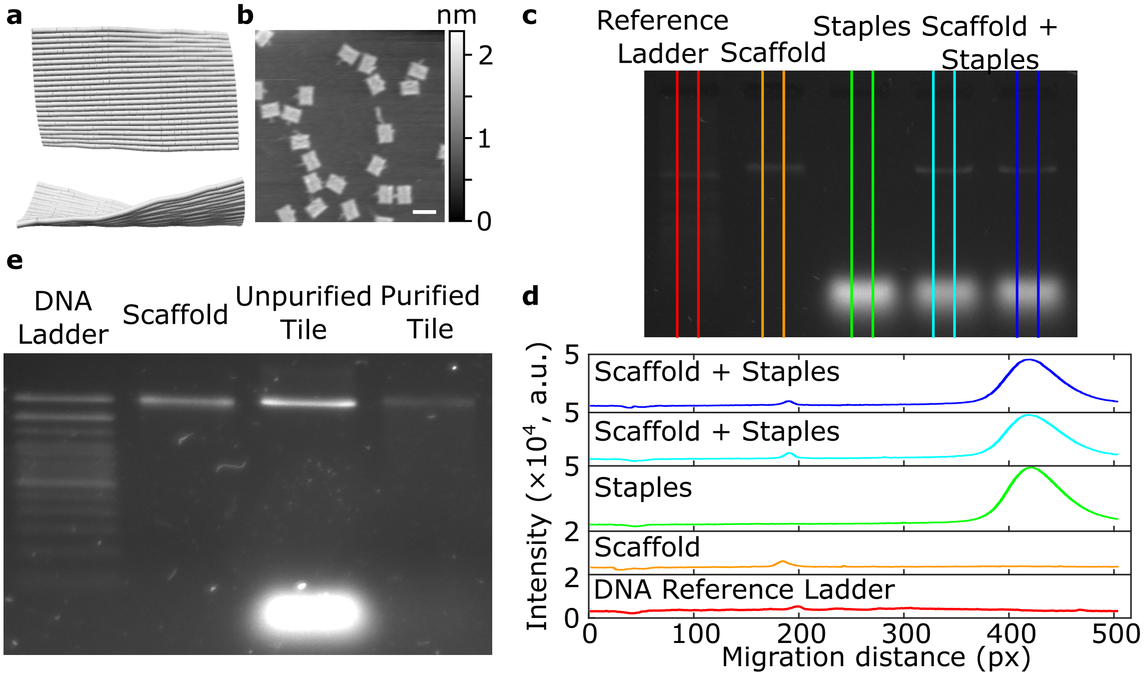


**Figure S1: The origami methodology allows to assemble DNA-based tiles with programmable functionality.** **a** Schematic representation of the top and side views of the Rothemund Rectangular Origami (RRO), as computed by Cando.[30,31] **b** Atomic force micrograph confirming the assembly of nanostructures with rectangular conformations. Scale bar = 100 nm. **c** Micrograph of agarose gel electrophoresis of scaffold, staples, and origami assembled from mixtures of scaffold and staples annealed with the temperature quenching protocol described in the Methods. **d** Intensity profiles of image in panel (b) showing the migration pattern of the scaffold, staples, and the folded origami. **e** Micrograph of agarose gel electrophoresis of scaffold alongside unpurified and purified DNA tiles following three cycles of PEG-induced precipitation as described in the Methods, showing the successful removal of excess staples.

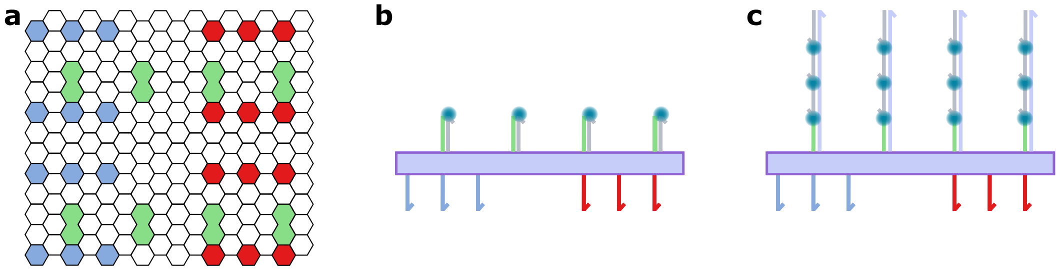


**Figure S2: Location of fluorescent beacons in DNA line-actants.** **a** Alexa488-conjugated oligonucleotides are incorporated to the RRO at 8 defined locations, highlighted in green in the hexagonal grid, by extending staples on their 5'-end to add docking domains, thus placing the fluorescent beacons on the face opposite to that hosting the anchor binding sites. **b** In the first design (used to collect z-stacks in Fig. 1 and Fig. 3 in the main text, we directly bind the fluorescent oligonucleotides (grey) to the docking domain (green), resulting in plates with ∼8 fluorophores. **c** Our second design, used in measurements for line-partitioning quantitation (Fig. 2), the docking domain (green) in turn binds an extender strand which bears 3× binding sites for the fluorescent oligonucleotide. In this case, each plate can feature up to 24× fluorophores.

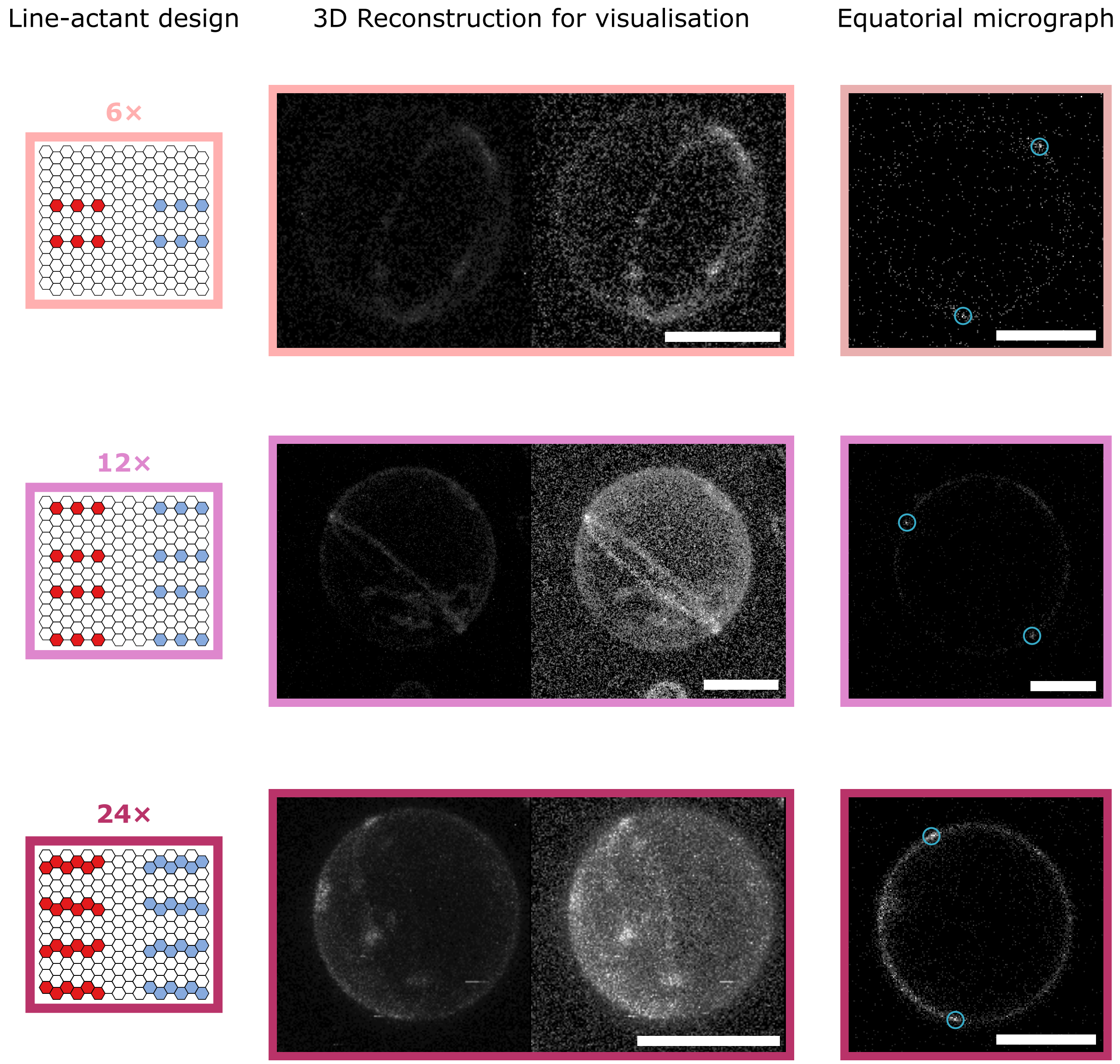


**Figure S3: DNA plates with rationally-positioned binding sites for multiple sT and dC anchor modules enrich the line-interface of de-mixed GUVs.** (Left) Schematic depictions of the tiles with the position of overhangs targeting sT (red) and dc (blue) anchoring modules. (Middle) 3D views of phase-separated vesicles with a Janus-like morphology showing line-accumulation of fluorescent (Alexa488) DNA plates with sets of either 6×, 12×, or 24× anchor-points, as reconstructed from confocal z-stacks using Volume Viewer (FIJI[32]) with (right) and without (left) contrast enhancement. (Right) Representative equatorial confocal micrographs of GUVs with line-actant plates, highlighting with blue circles the location of the line-interfaces, which were used for quantitation of line-accumulation (see Fig. 2 in the main text). Scale bars = 10 $\mu$m.

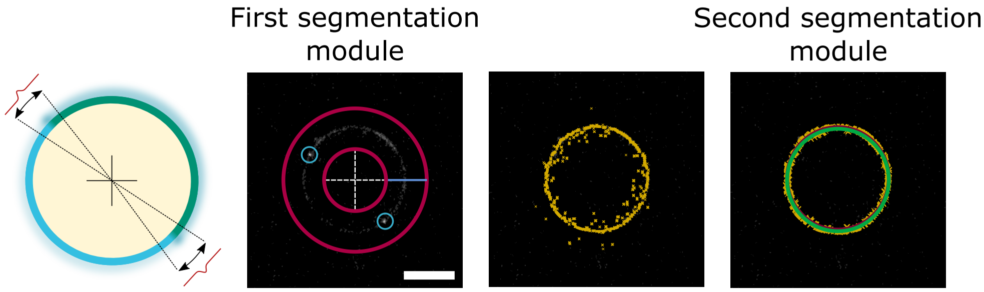


**Figure S4: Micrograph segmentation allows to reconstruct the interfacial fluorescence intensity profile.** The analysis pipeline operates on equatorial micrographs, recording the signal of Alexa488 fluorophores on the DNA origami. The custom-built script loads the micrograph, where the user identifies rough locations for three positions: i) the boundaries between lipid phases, and ii) a location of the $L_\mathrm{d}$ domain. Using these positions, a circle is fitted as reference so to enclose the membrane within a ring. Image processing proceeds with a first segmentation module, where a line connecting the two concentric circles (in red) is projected radially using polar coordinates. Thus, radial intensity profiles are acquired, from $\theta = 0$ to $2\pi$ in steps of $\pi/900$. From individual radial profiles, we extracted the location of the membrane *via* numerical integration around the membrane peak, as determined with a Gaussian fit. To further account for imperfect segmentation due to noise, these positions (shown in yellow in the middle micrograph) are filtered in a second segmentation module. The latter uses a subset of the membrane positions, namely those that are within $\pm 5\%$ of the estimated radius of the vesicle. These points are then subjected to a circle fitting routine for each lipid (quasi-hemispherical) domain, and the average membrane intensity values are extracted at a distance equal to the radius of the circle (in red and green in the right micrograph). These points are used to construct an intensity profile around the location of the line-interface, as described in Supplementary Note II. Scale bar = $10\,\mu$m.

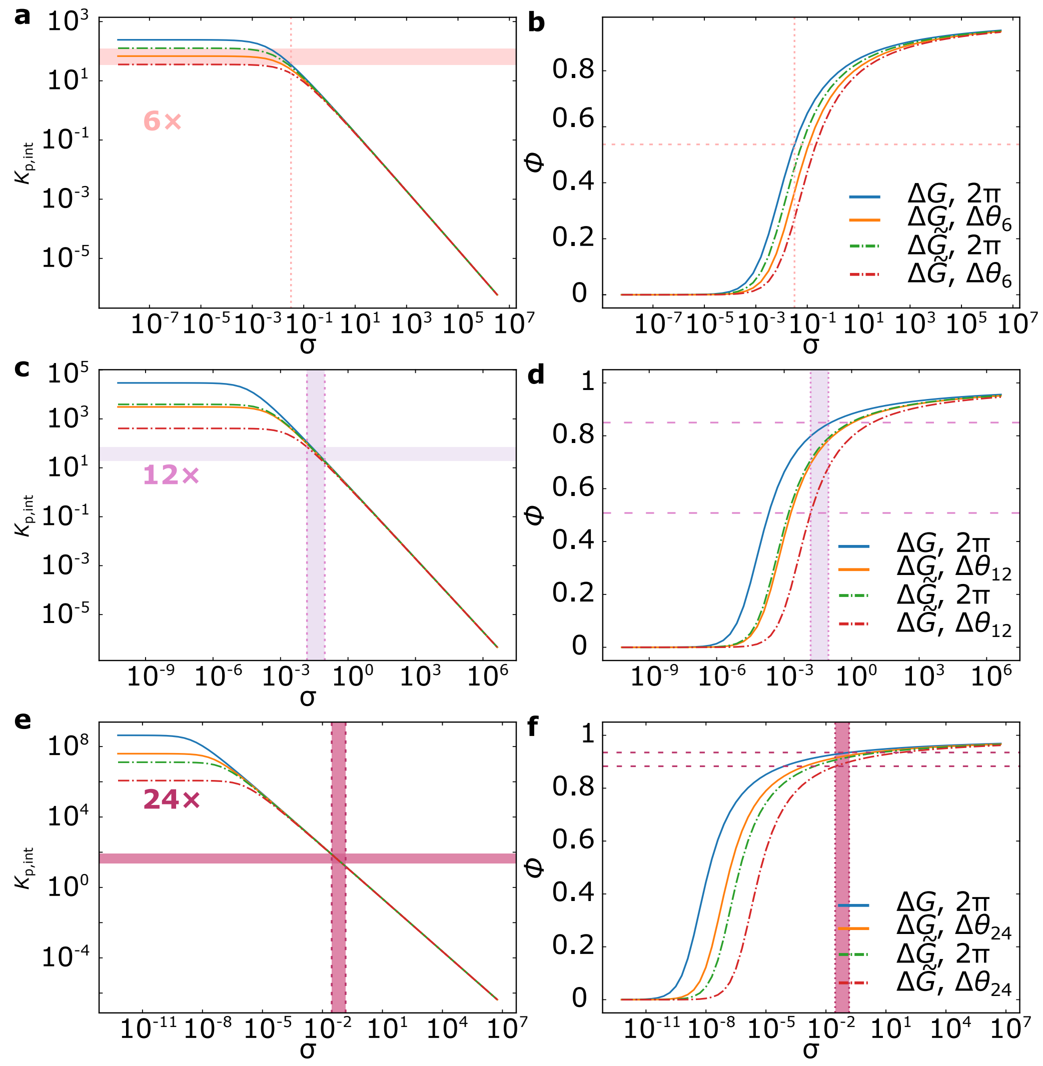


**Figure S5: Comparing experimental trends with our numerical model reveals the regulatory effect on line-partitioning afforded by DNA nanostructures. a–c** Plots of the partitioning coefficient ($K_{\mathrm{p,int}}$) as a function of surface coverage ($\sigma$), computed using $\xi = 8\,\mathrm{nm}$ and the parameter values compiled in Table S6 for: i) the free-energy contributions for partitioning assuming all anchor-points are present in the line-actant ($\Delta G$, solid lines); or ii) the free-energy contributions for partitioning correcting for the probability of staple incorporation ($\Delta \tilde{G}$, dash-dot lines), as quantified by Jungmann and co-workers; iii) the reduced rotational freedom of the tiles upon accumulating at the line-interface due to the position of the anchor-points, which we describe with $\Delta\theta_{\mathrm{n}}$ (orange/red curves); and finally iv) tiles with full rotational freedom ($\Delta\theta = 2\pi$, blue/green curves). By intersecting the experimental line-partitioning coefficients ± standard deviation ($K_{\mathrm{p,int}} \pm \delta K_{\mathrm{p,int}}$, horizontal shaded region) with the curves from the theoretical predictions, we can infer surface coverages ($\sigma$) relevant to the scenarios outlined above (i–iv), providing us with a degree of uncertainty (vertical shaded region). **d–f** We then apply the inferred ranges of $\sigma$ to estimate the fraction of occupied line-interface ($\phi$), computed by solving Eq. S14 for scenarios i–iv, which provides us with lower and upper bounds for $\phi_{\mathrm{n}\times}$. The resulting ranges of $\sigma$ and $\phi$ are summarised in Fig. S6 and Fig. 2 (main text), respectively.

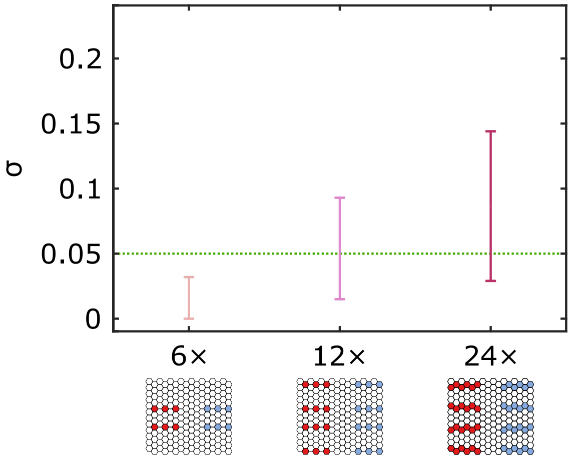


**Figure S6: The predicted surface coverage is similar to the expected experimental value** As discussed in Supplementary Note III, we inferred the surface coverage ($\sigma = \rho L_1 L_2$) of line-actants on membranes in view of their experimental line-partitioning tendencies (Fig. 2 in the main text and Supplementary Fig. S5. The estimated ranges of $\sigma$ for each line-actant design is similar to the anticipated experimental values ($\sigma_{\text{exp}} \sim 0.05$, green dashed line). In the case of the 6× design, we observe ranges of surface coverage slightly below the nominal experimental value, which we ascribe to the lower membrane-affinity of the nanostructures given the fewer number of anchor-points they feature.

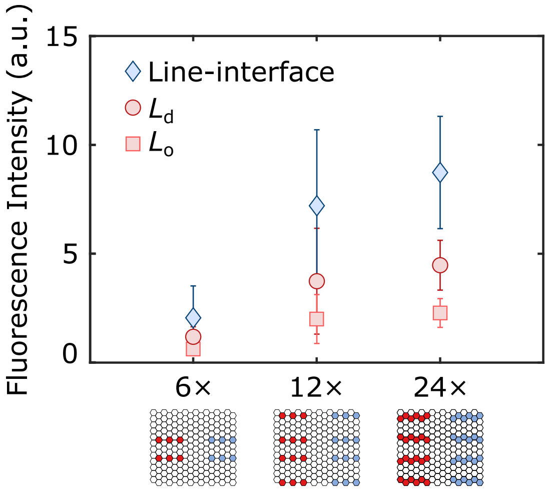


**Figure S7: Lower number of anchors results in a lower surface density of line-actants.** The absolute fluorescence intensities quantified from equatorial confocal micrographs show that nanostructures equipped with 6× have systematically lower membrane fluorescence intensities than those detected for GUVs decorated with 12× and 24× dC and sT anchor-points, thus suggesting lower surface densities. The latter can be due to an overall lower affinity of the nano-devices for the membrane.

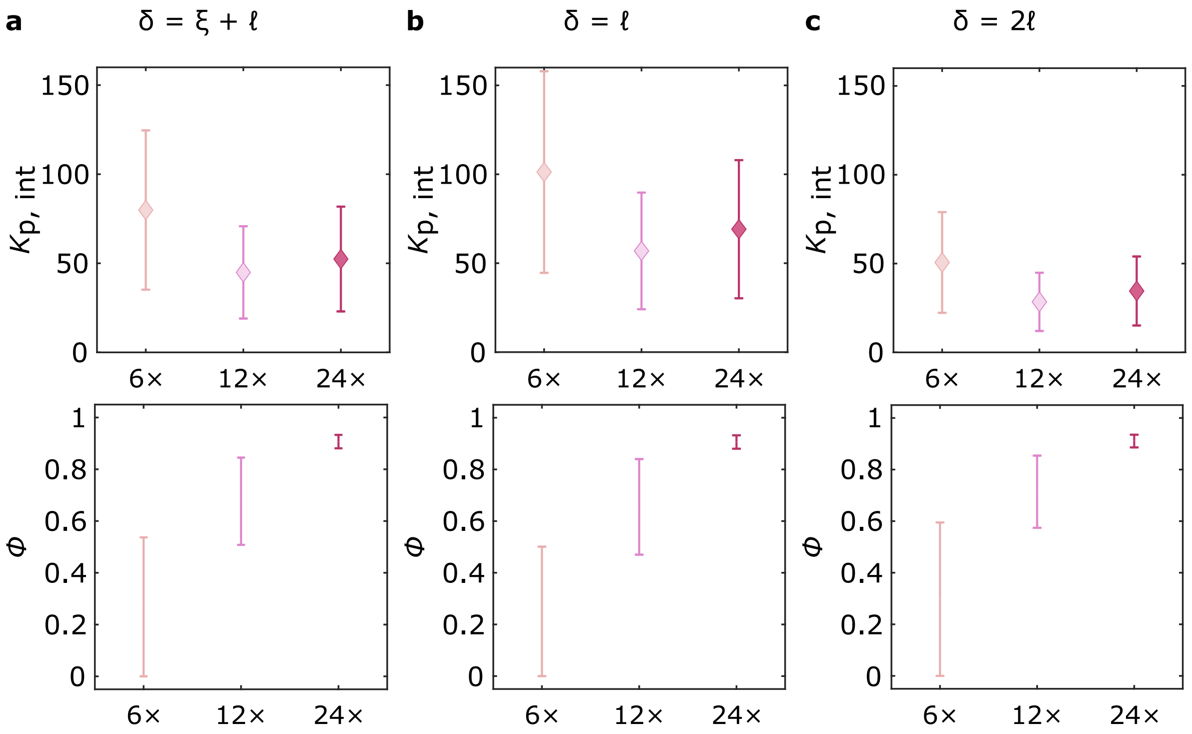


**Figure S8: Error propagation of $\xi$ to line-accumulation observables.** The uncertainty on the estimation of $\xi$ propagates to the values of $K_{\mathrm{p,int}}$, while having a negligible influence on the estimation of $\phi$. Comparing $\xi = 8\,\mathrm{nm}$ (panel **a**) with both lower (panel **b**, $\xi = 0$) and upper (panel **c**, $\xi = 30\,\mathrm{nm}$, for 6× and 12× tiles, or $\xi = 25\,\mathrm{nm}$ for 24× tiles) physical bounds on the thickness of the line-interface shows that the values of $K_{\mathrm{p,int}}$ shift slightly, and are all within a factor of two from each other. Conversely, the values of $\phi$ only negligibly change, and preserve the trend of increased line-accumulation with increasing number of anchor-points on the line-actants.

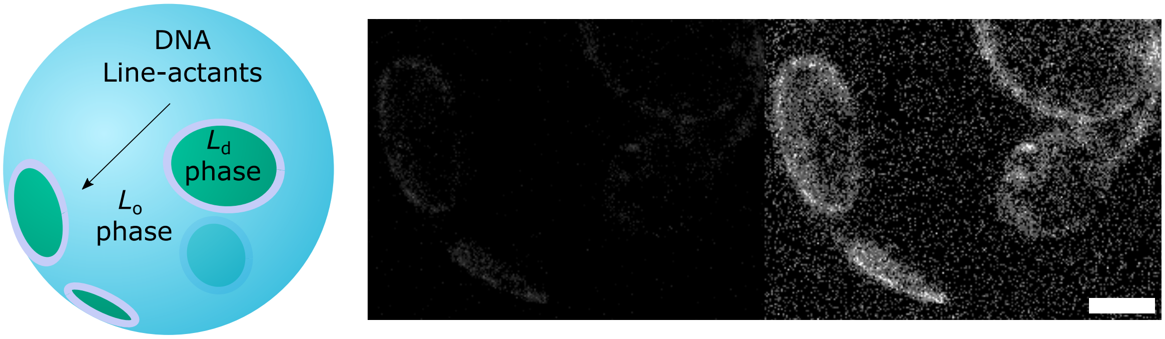


**Figure S9: Line-active DNA nanostructures stabilise lipid domains by accumulating at line-interfaces.** Line-accumulation of DNA nanostructures scaffolds lipid domains and renders them stable against coalescence, as shown with a 3D view of a reconstructed phase-separated vesicle after quenching below the miscibility transition temperature. Line-accumulation of fluorescent (Alexa488) DNA plates is readily noticeable, showing stable domains rendered from a confocal z-stack using Volume Viewer (FIJI[32]) with (right) and without (left) contrast enhancement. Scale bar = 10 $\mu$m.

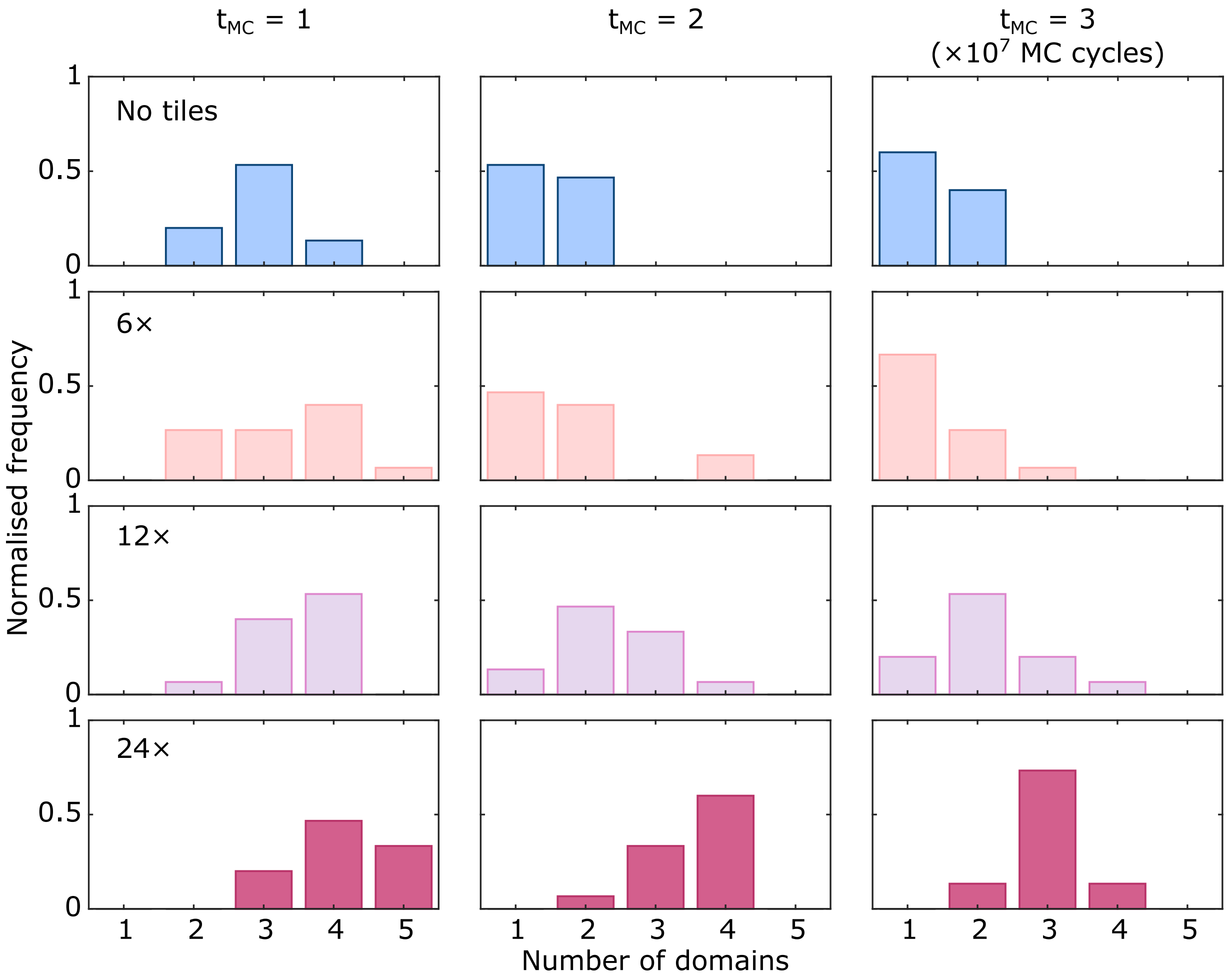


**Figure S10: Time-evolution of domain stabilisation with line-active tiles.** Frequency histograms comparing the number of domains in simulation runs, normalised to the total number of trajectories, at various $t_{\text{MC}}$ in the presence of line-actant designs 6× ($n = 15$ trajectories), 12× ($n = 15$ trajectories), or 24× ($n = 15$ trajectories), comparing them to simulations run without DNA tiles ($n = 15$ trajectories).

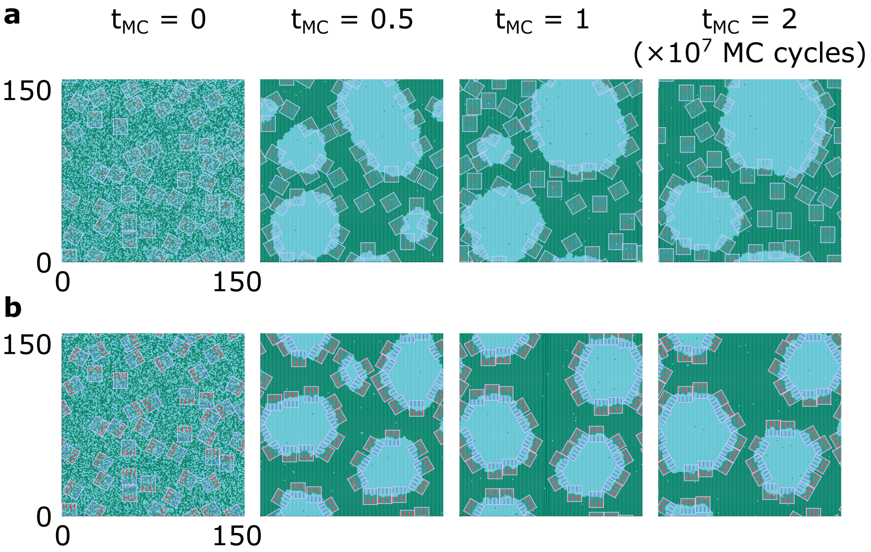


**Figure S11: Temporal evolution of domain stabilisation with line-actant variants.** Domain stabilisation demonstrated with representative snapshots at various $t_{\mathrm{MC}}$ for simulations with: **a** 6× or **b** 24× DOLAs.

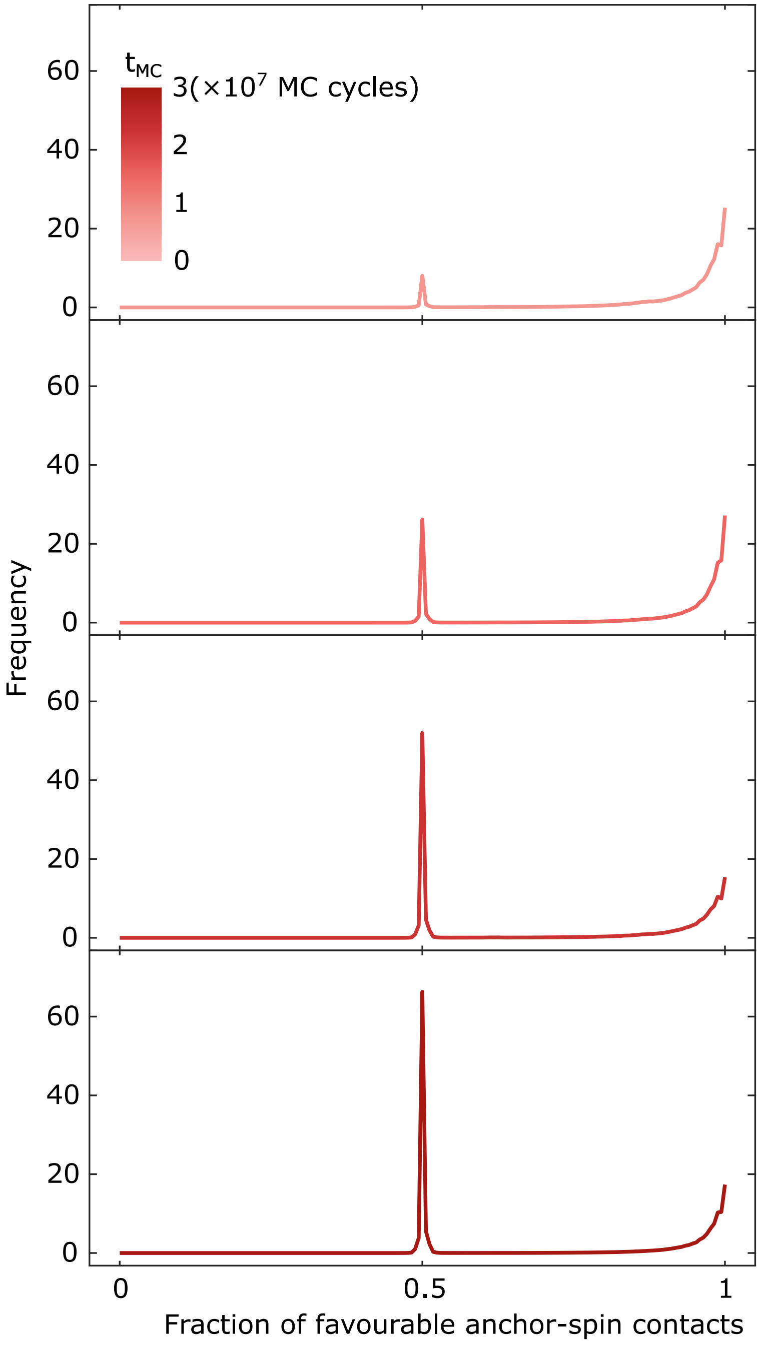


**Figure S12: The fraction of favourable anchor-spin contacts are used to assess line-accumulation of DNA tiles.** In order to compute the number of tiles adsorbed at the line-interface, we use the fraction of favourable anchor-spin contacts. A tile is said to be at the line when featuring at least 80% of favourable anchor-spin interactions relative to the total contacts. At early $t_{\mathrm{MC}}$, when most tiles are adsorbed at the line-interface, we observe a high frequency of favourable anchor-spin contacts $\sim 1$, with only few occurrences of 0.5 contacts. Late $t_{\mathrm{MC}}$ indicate line-interface minimisation and the associated translocation of the tiles from the interface to the green phase, which results in half of the anchor-points being hosted in their unfavourable phase. The latter thus shifts the frequencies of favourable anchor-spin contacts to 0.5, increasing the height of the peak.

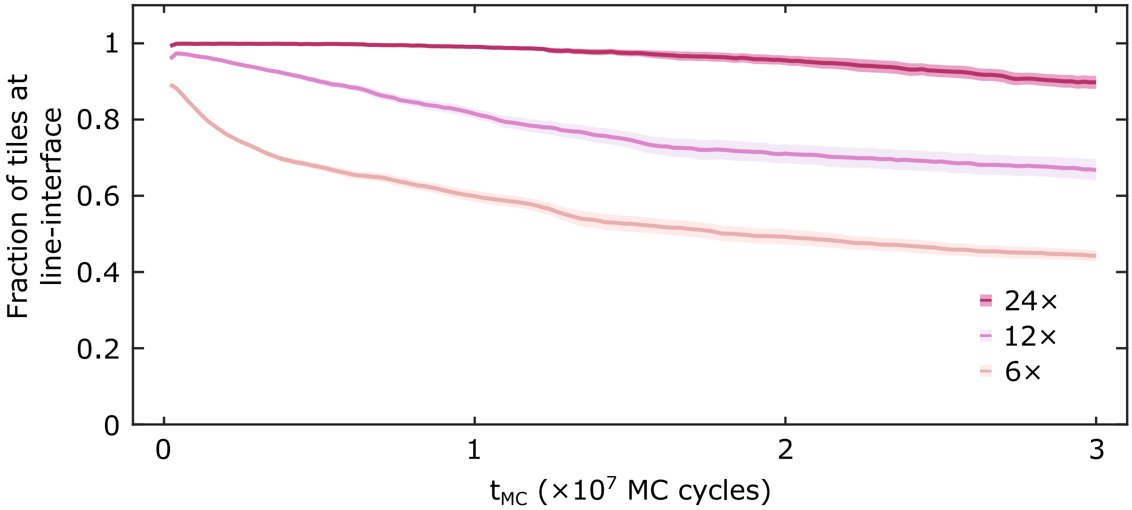


**Figure S13: Domain coarsening drives tile desorption from line-interfaces.** As the simulations progress, the fraction of tiles adsorbed at line-interfaces decreases to levels that increase monotonically with the number of anchor-points. Note that, even for the line-accumulating design with the most favourable free-energy gain (24×), there is still a substantial decrease, hinting at the prevalence of Ostwald ripening.

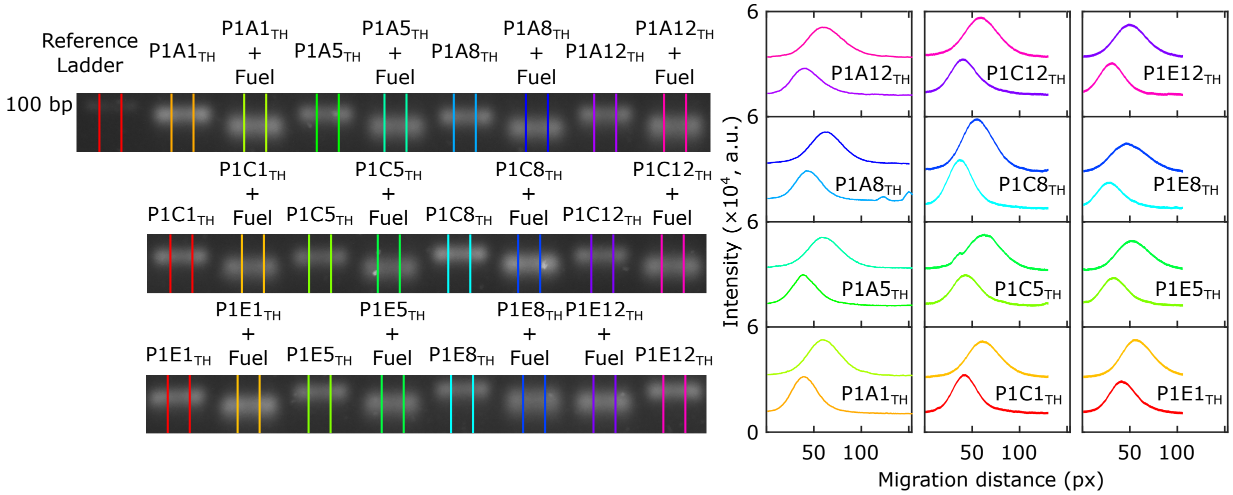


**Figure S14: Reconfigurability of anchor-points through toeholding reactions confirmed with AGE.** Agarose gel electrophoresis demonstrates functionality of the toeholding reaction in each (dC) anchor-point used for the dynamic line-actants in Fig. 5 and Fig. 6 in the main text. Each dC-targeting staple, bearing the toehold with sequence $\alpha$, was assembled via annealing with their corresponding (non-cholesterolised) dC anchor module. Gel electrophoresis was run on these nanostructures, and also in the presence of Fuel, as indicated in each lane. The toeholding reaction catalysed by the Fuel detaches the staple from the anchor-modules, resulting in smaller constructs and thus different migration profiles. The shift and broadening of the peaks after the addition of Fuel in the intensity profiles are indicative of construct reconfiguration into smaller devices, thus confirming the molecular response for each staple.

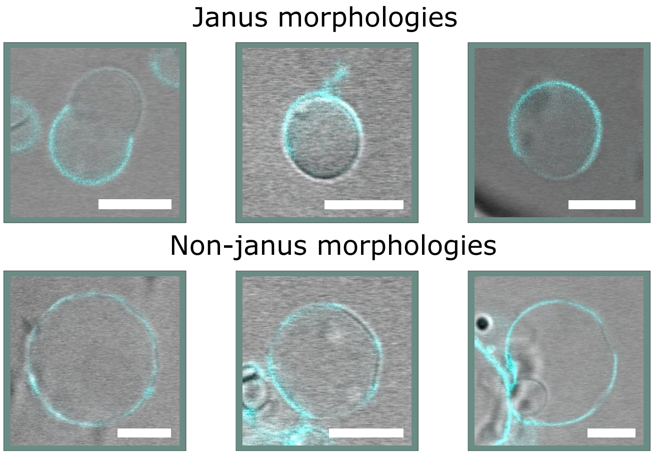


**Figure S15: Changes in spontaneous curvature in GUVs can stabilise lipid domains.** Confocal micrograph overlays showing representative examples of Janus-looking line-actant functionalised GUVs, after a heating-cooling program and exposed to Fuel, coexisting with vesicles that still feature Non-janus morphologies. In the latter case, it is noticeable that the spontaneous curvature of the vesicle has deviated from that of a sphere, likely due to small osmotic imbalances due to the addition of Fuel, which allowed lipid domains to retain stability against coalescence. In cyan, the fluorescent signal of the DOLAs, fluorescently labelled with Alexa488. All scale bars = 10 $\mu$m.

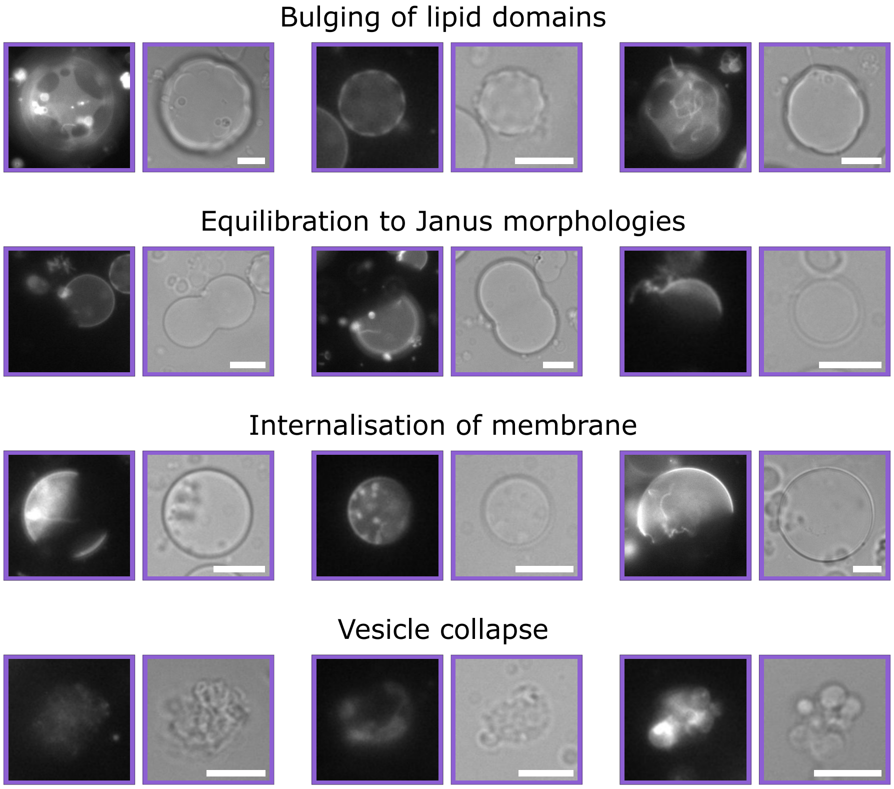


**Figure S16: Influence of hyperosmosis in GUVs.** Inducing hyperosmosis at $C_\mathrm{O}/C_\mathrm{I} \approx 1.28$ leads to sample heterogeneity across vesicles. We observed four possible outcomes in hyperosmolar conditions: i) bulging of line-actant stabilised lipid domains; ii) equilibration to Janus-like morphologies; iii) uptake of membrane to reduce the excess surface area, leading to taut vesicles, and; iv) vesicle collapse, likely due to the fast changes in osmotic conditions. All scale bars = $10\,\mu$m.

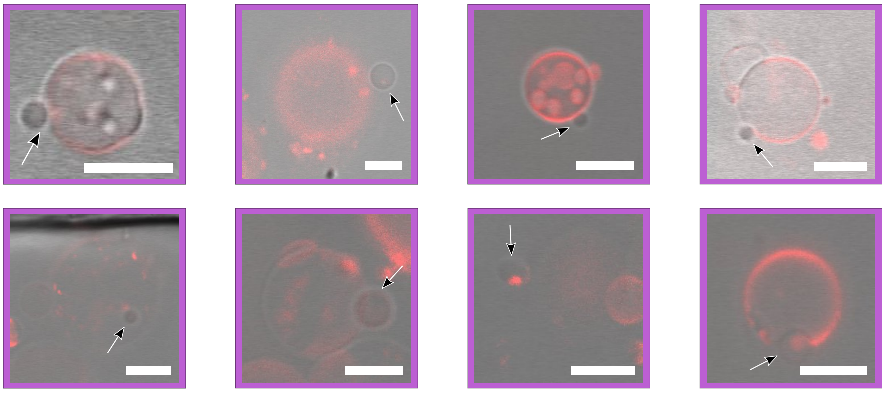

**Figure S17: Fission in line-actant functionalised liposomes.** Confocal micrograph overlays of domain fission events, including those in Fig. 6 in the main text, where GUVs featuring DOLAs were exposed to hyperosmotic shock ($C_{\mathrm{O}}/C_{\mathrm{I}} \approx 1.28$) and supplemented with Fuel to trigger line-actant reconfiguration. Vesicles immersed in buffer with Fuel mostly featured the expected Janus-like morphologies, suggesting domain coarsening. In these conditions, we identified a further 7 events, shown with representative examples here alongside those included in Fig. 6, indicating that fission occurred in some synthetic cells, while other vesicles fully relaxed into Janus morphologies. Both the competition between line energy and fluctuation entropy as well as the kinetics of toehold-induced domain re-organisation could account for sample heterogeneity. Specifically, unlike measurements in Fig. 5, the progression of strand displacement is not limited by the diffusion of Fuel molecules and may occur in timescales comparable to those of osmosis, which could in some instances lead to equilibration to single-domain phases prior to domain budding, at which point fission can no longer proceed. The red signal detected is that of TexasRed-DHPE, which stains the $L_{\mathrm{d}}$ phase. All scale bars = $10\,\mu$m.

### Supplementary Movies Key

**Supplementary Movie 1**: Representative simulation trajectory with 12× DNA-origami Line-actants following the system after quenching for $2 \times 10^7$ MC cycles. Box size: 320×372 lattice points.

**Supplementary Movie 2**: Representative simulation trajectory with 12× DNA-origami Line-actants following the system after quenching for $3 \times 10^7$ MC cycles. Box size: 160×172 lattice points.

**Supplementary Movie 3**: Representative simulation trajectory lacking DNA-origami Line-actants following the system after quenching for $3 \times 10^7$ MC cycles. Box size: 160×172 lattice points.

**Supplementary Movie 4**: Representative simulation trajectory with 24× DNA-origami Line-actants following the system after quenching for $3 \times 10^7$ MC cycles. Box size: 160×172 lattice points.

**Supplementary Movie 5**: Representative simulation trajectory with 6× DNA-origami Line-actants following the system after quenching for $3 \times 10^7$ MC cycles. Box size: 160×172 lattice points.

**Supplementary Movie 6**: Evolution of $L_o$-domain fission event in a representative vesicle observed with brightfield and epi-fluorescence microscopy. Time-stamps refer to the time elapsed after initial acquisition. Fuel was added 2 minutes before acquisition began. The fluorescent marker is TexasRed-DHPE, which labels the $L_d$ phase. Scale bar: $10\,\mu$m.

**Supplementary Movie 7**: Evolution of $L_o$-domain fission event in a representative vesicle observed with overlaid confocal and brightfield microscopy. Time-stamps refer to the time elapsed after initial acquisition. Fuel was added 90 minutes before acquisition began. The fluorescent marker is TexasRed-DHPE, which labels the $L_{\mathrm{d}}$ phase. Scale bar: 10 $\mu$m.

### Supplementary Tables

Table S1: **Staple sequences for DNA origami tile.** Sequences of oligonucleotides comprising the origami plate, highlighting in **bold** the overhangs extended in selected staples to accommodate domains to bind anchor-modules dC (blue) and sT (red) as well as the fluorescent beacon $\mathrm{Lin}^{f,*}$ or Extender-3×$\mathrm{Lin}^{f,*}$ (green).

| Name | Sequence |
|---|---|
| P1A01 | TTTTCACTCAAAGGGCGAAAAACCATCACC**GGTTTGTTGTTGTGTTGG** |
| P1A02 | GTCGACTTCGGCCAACGCGCGGGGTTTTTC |
| P1A03 | TGCATCTTTCCCAGTCACGACGGCCTGCAG |
| P1A04 | TAATCAGCGGATTGACCGTAATCGTAACCG |
| P1A05 | AACGCAAAATCGATGAACGGTACCGGTTGA**GGTTTGTTGTTGTGTTGG** |
| P1A06 | AACAGTTTTGTACCAAAAACATTTTATTTC |
| P1A07 | TTTACCCCAACATGTTTTAAATTTCCATAT |
| P1A08 | TTTAGGACAAATGCTTTAAACAATCAGGTC**GGTTTGTTGTTGTGTTGG** |
| P1A09 | CATCAAGTAAAACGAACTAACGAGTTGAGA |
| P1A10 | AATACGTTTGAAAGAGGACAGACTGACCTT |
| P1A11 | AGGCTCCAGAGGCTTTGAGGACACGGGTAA |
| P1A12 | AGAAAGGAACAACTAAAGGAATTCAAAAAAA**GGTTTGTTGTTGTGTTGG** |
| P1B01 | CAAATCAAGTTTTTTGGGGTCGAAACGTGGA**GGTTTGTTGTTGTGTTGG** |
| P1B02 | CTCCAACGCAGTGAGACGGGCAACCAGCTGCA |
| P1B03 | TTAATGAACTAGAGGATCCCCGGGGGGTAACG |
| P1B04 | CCAGGGTTGCCAGTTTGAGGGGACCCGTGGGA**GGTTTGTTGTTGTGTTGG** |
| P1B05 | ACAAACGGAAAAGCCCCAAAAACACTGGAGCA |
| P1B06 | AACAAGAGGGATAAAAATTTTTAGCATAAAGC |
| P1B07 | TAAATCGGGATTCCCAATTCTGCGATATAATG |
| P1B08 | CTGTAGCTTGACTATTATAGTCAGTTCATTGA**GGTTTGTTGTTGTGTTGG** |
| P1B09 | ATCCCCCTATACCACATTCAACTAGAAAAATC |
| P1B10 | TACGTTAAAGTAATCTTGACAAGAACCGAACT |
| P1B11 | GACCAACTAATGCCACTACGAAGGGGGTAGCA**GGTTTGTTGTTGTGTTGG** |
| P1B12 | ACGGCTACAAAAGGAGCCTTTAATGTGAGAAT |
| P1C01 | AGCTGATTGCCCTTCAGAGTCCACTATTAAAGGGTGCCGT**GGTTTGTTGTTGTGTTGG** |
| P1C04 | GTATAAGCCAACCCGTCGGATTCTGACGACAGTATCGGCCGCAAGGCG |
| P1C05 | TATATTTTGTCATTGCCTGAGAGTGGAAGATT**GGTTTGTTGTTGTGTTGG** |
| P1C06 | GATTTAGTCAATAAAGCCTCAGAGAACCCTCA |
| P1C07 | CGGATTGCAGAGCTTAATTGCTGAAACGAGTA |
| P1C08 | ATGCAGATACATAACGGGAATCGTCATAAATAAAGCAAAG**GGTTTGTTGTTGTGTTGG** |
| P1C11 | TTTATCAGGACAGCATCGGAACGACACCAACCTAAAACGAGGTCAATC |
| P1C12 | ACAACTTTCAACAGTTTCAGCGGATGTATCGG**GGTTTGTTGTTGTGTTGG** |
| P1D01 | AAAGCACTAAATCGGAACCCTAATCCAGTT**GGTTTGTTGTTGTGTTGG** |

| | |
|---|---|
| P1D02 | TGGAACAACCGCCTGGCCCTGAGGCCCGCT |
| P1D03 | TTCCAGTCGTAATCATGGTCATAAAAGGGG |
| P1D04 | GATGTGCTTCAGGAAGATCGCACAATGTGA**GGTTTGTTGTTGTGTTGG** |
| P1D05 | GCGAGTAAAAATATTTAAATTGTTACAAAG |
| P1D06 | GCTATCAGAAATGCAATGCCTGAATTAGCA |
| P1D07 | AAATTAAGTTGACCATTAGATACTTTTGCG |
| P1D08 | GATGGCTTATCAAAAAGATTAAGAGCGTCC**GGTTTGTTGTTGTGTTGG** |
| P1D09 | AATACTGCCCAAAAGGAATTACGTGGCTCA |
| P1D10 | TTATACCACCAAATCAACGTAACGAACGAG |
| P1D11 | GCGCAGACAAGAGGCAAAAGAATCCCTCAG**GGTTTGTTGTTGTGTTGG** |
| P1D12 | CAGCGAAACTTGCTTTCGAGGTGTTGCTAA |
| P1E01 | AGCAAGCGTAGGGTTGAGTGTTGTAGGGAGCC**GGTTTGTTGTTGTGTTGG** |
| P1E02 | CTGTGTGATTGCGTTGCGCTCACTAGAGTTGC |
| P1E03 | GCTTTCCGATTACGCCAGCTGGCGGCTGTTTC |
| P1E04 | ATATTTTGGCTTTCATCAACATTATCCAGCCA |
| P1E05 | TAGGTAAACTATTTTTGAGAGATCAAACGTTA**GGTTTGTTGTTGTGTTGG** |
| P1E06 | AATGGTCAACAGGCAAGGCAAAGAGTAATGTG |
| P1E07 | CGAAAGACTTTGATAAGAGGTCATATTTCGCA |
| P1E08 | TAAGAGCAAATGTTTAGACTGGATAGGAAGCC**GGTTTGTTGTTGTGTTGG** |
| P1E09 | TCATTCAGATGCGATTTTAAGAACAGGCATAG |
| P1E10 | ACACTCATCCATGTTACTTAGCCGAAAGCTGC |
| P1E11 | AAACAGCTTTTTGCGGGATCGTCAACACTAAA |
| P1E12 | TAAATGAATTTTCTGTATGGGATTAATTTCTT**GGTTTGTTGTTGTGTTGG** |
| P1F01 | CCCGATTTAGAGCTTGACGGGGAAAAAGAATA**GGTTTGTTGTTGTGTTGG** |
| P1F02 | GCCCGAGAGTCCACGCTGGTTTGCAGCTAACT |
| P1F03 | CACATTAAAATTGTTATCCGCTCATGCGGGCC |
| P1F04 | TCTTCGCTGCACCGCTTCTGGTGCGGCCTTCC**GGTTTGTTGTTGTGTTGG** |
| P1F05 | TGTAGCCATTAAAATTCGCATTAAATGCCGGA |
| P1F06 | GAGGGTAGGATTCAAAAGGGTGAGACATCCAA |
| P1F07 | TAAATCATATAACCTGTTTAGCTAACCTTTAA |
| P1F08 | TTGCTCCTTTCAAATATCGCGTTTGAGGGGGT**GGTTTGTTGTTGTGTTGG** |
| P1F09 | AATAGTAAACACTATCATAACCCTCATTGTGA |
| P1F10 | ATTACCTTTGAATAAGGCTTGCCCAAATCCGC |
| P1F11 | GACCTGCTCTTTGACCCCCAGCGAGGGAGTTA**GGTTTGTTGTTGTGTTGG** |
| P1F12 | AAGGCCGCTGATACCGATAGTTGCGACGTTAG |
| P1G01 | CCCAGCAGGCGAAAAATCCCTTATAAATCAAGCCGGCG |
| P1G04 | TAAATCAAAATAATTCGCGTCTCGGAAACCAGGCAAAGGGAAGG |
| P1G05 | GAGACAGCTAGCTGATAAATTAATTTTTGT |
| P1G06 | TTTGGGGATAGTAGTAGCATTAAAAGGCCG |
| P1G07 | GCTTCAATCAGGATTAGAGAGTTATTTTCA |
| P1G08 | CGTTTACCAGACGACAAAGAAGTTTTGCCATAATTCGA |

| | |
|---|---|
| P1G11 | TGACAACTCGCTGAGGCTTGCATTATACCAAGCGCGATGATAAA |
| P1G12 | TCTAAAGTTTTGTCGTCTTTCCAGCCGACAA |
| P1H01 | TCAATATCGAACCTCAAATATCAATTCCGAAA |
| P1H02 | GCAATTCACATATTCCTGATTATCAAAGTGTA |
| P1H03 | AGAAAACAAAGAAGATGATGAAACAGGCTGCG |
| P1H04 | ATCGCAAGTATGTAAATGCTGATGATAGGAAC |
| P1H05 | GTAATAAGTTAGGCAGAGGCATTTATGATATT |
| P1H06 | CCAATAGCTCATCGTAGGAATCATGGCATCAA |
| P1H07 | AGAGAGAAAAAAATGAAAATAGCAAGCAAACT |
| P1H08 | TTATTACGAAGAACTGGCATGATTGCGAGAGG |
| P1H09 | GCAAGGCCTCACCAGTAGCACCATGGGCTTGA |
| P1H10 | TTGACAGGCCACCACCAGAGCCGCGATTTGTA |
| P1H11 | TTAGGATTGGCTGAGACTCCTCAATAACCGAT |
| P1H12 | TCCACAGACAGCCCTCATAGTTAGCGTAACGA |
| P2A01 | AACGTGGCGAGAAAGGAAGGGAAACCAGTAA |
| P2A02 | TCGGCAAATCCTGTTTGATGGTGGACCCTCAA |
| P2A03 | AAGCCTGGTACGAGCCGGAAGCATAGATGATG |
| P2A04 | CAACTGTTGCGCCATTCGCCATTCAAACATCA |
| P2A05 | GCCATCAAGCTCATTTTTTAACCACAAATCCA |
| P2A06 | CAACCGTTTCAAATCACCATCAATTCGAGCCA |
| P2A07 | TTCTACTACGCGAGCTGAAAAGGTTACCGCGC |
| P2A08 | CCAACAGGAGCGAACCAGACCGGAGCCTTTAC |
| P2A09 | CTTTTGCAGATAAAAACCAAAATAAAGACTCC |
| P2A10 | GATGGTTTGAACGAGTAGTAAATTTACCATTA |
| P2A11 | TCATCGCCAACAAAGTACAACGGACGCCAGCA |
| P2A12 | ATATTCGGAACCATCGCCCACGCAGAGAAGGA |
| P2B01 | TAAAAGGGACATTCTGGCCAACAAAGCATC |
| P2B02 | ACCTTGCTTGGTCAGTTGGCAAAGAGCGGA |
| P2B03 | ATTATCATTCAATATAATCCTGACAATTAC |
| P2B04 | CTGAGCAAAAATTAATTACATTTTGGGTTA |
| P2B05 | TATAACTAACAAAGAACGCGAGAACGCCAA |
| P2B06 | CATGTAATAGAATATAAAGTACCAAGCCGT |
| P2B07 | TTTTATTTAAGCAAATCAGATATTTTTTGT |
| P2B08 | TTAACGTCTAACATAAAAACAGGTAACGGA |
| P2B09 | ATACCCAACAGTATGTTAGCAAATTAGAGC |
| P2B10 | CAGCAAAAGGAAACGTCACCAATGAGCCGC |
| P2B11 | CACCAGAAAGGTTGAGGCAGGTCATGAAAG |
| P2B12 | TATTAAGAAGCGGGGTTTTGCTCGTAGCAT |
| P2C01 | TCAACAGTTGAAAGGAGCAAATGAAAAATCTAGAGATAGA**GGTGTAGTTAATGGGAGT** |
| P2C02 | XXXXXXXXXXXXXXXXATTAAGTTTACCGAGCTCGAATTCGGGAAACCTGTCGTGC |
| P2C03 | XXXXXXXXXXXXXXXXATAAGGGAACCGGATATTCATTACGTCAGGACGTTGGGAA |

| | |
|---|---|
| P2C04 | TCAAATATAACCTCCGGCTTAGGTAACAATTTCATTTGAAGGCGAATT |
| P2C05 | GTAAAGTAATCGCCATATTTAACAAAACTTTT**GGTGTAGTTAATGGGAGT** |
| P2C06 | TATCCGGTCTCATCGAGAACAAGCGACAAAAG |
| P2C07 | TTAGACGGCCAAATAAGAAACGATAGAAGGCT |
| P2C08 | CGTAGAAAATACATACCGAGGAAACGCAATAAGAAGCGCA**GGTGTAGTTAATGGGAGT** |
| P2C09 | XXXXXXXXXXXXXXXXGCGATCGGCAATTCCACACAACAGGTGCCTAATGAGTG |
| P2C10 | XXXXXXXXXXXXXXXXTTGTGTCGTGACGAGAAACACCAAATTTCAACTTTAAT |
| P2C11 | GCGGATAACCTATTATTCTGAAACAGACGATTGGCCTTGAAGAGCCAC |
| P2C12 | TCACCAGTACAAACTACAACGCCTAGTACCAG**GGTGTAGTTAATGGGAGT** |
| P2D01 | ACCCTTCTGACCTGAAAGCGTAAGACGCTGAG**GGTGTAGTTAATGGGAGT** |
| P2D02 | AGCCAGCAATTGAGGAAGGTTATCATCATTTT |
| P2D03 | GCGGAACATCTGAATAATGGAAGGTACAAAAT |
| P2D04 | CGCGCAGATTACCTTTTTTAATGGGAGAGACT**GGTGTAGTTAATGGGAGT** |
| P2D05 | ACCTTTTTATTTTAGTTAATTTCATAGGGCTT |
| P2D06 | AATTGAGAATTCTGTCCAGACGACTAAACCAA |
| P2D07 | GTACCGCAATTCTAAGAACGCGAGTATTATTT |
| P2D08 | ATCCCAATGAGAATTAACTGAACAGTTACCAG**GGTGTAGTTAATGGGAGT** |
| P2D09 | AAGGAAACATAAAGGTGGCAACATTATCACCG |
| P2D10 | TCACCGACGCACCGTAATCAGTAGCAGAACCG |
| P2D11 | CCACCCTCTATTCACAAACAAATACCTGCCTA**GGTGTAGTTAATGGGAGT** |
| P2D12 | TTTCGGAAGTGCCGTCGAGAGGGTGAGTTTCG |
| P2E01 | CTTTAGGGCCTGCAACAGTGCCAATACGTG**GGTGTAGTTAATGGGAGT** |
| P2E02 | CTACCATAGTTTGAGTAACATTTAAAATAT |
| P2E03 | CATAAATCTTTGAATACCAAGTGTTAGAAC |
| P2E04 | CCTAAATCAAAATCATAGGTCTAAACAGTA |
| P2E05 | ACAACATGCCAACGCTCAACAGTCTTCTGA**GGTGTAGTTAATGGGAGT** |
| P2E06 | GCGAACCTCCAAGAACGGGTATGACAATAA |
| P2E07 | AAAGTCACAAAATAAACAGCCAGCGTTTTA |
| P2E08 | AACGCAAAGATAGCCGAACAAACCCTGAAC**GGTGTAGTTAATGGGAGT** |
| P2E09 | TCAAGTTTCATTAAAGGTGAATATAAAAGA |
| P2E10 | TTAAAGCCAGAGCCGCCACCCTCGACAGAA |
| P2E11 | GTATAGCAAACAGTTAATGCCCAATCCTCA |
| P2E12 | AGGAACCCATGTACCGTAACACTTGATATAA**GGTGTAGTTAATGGGAGT** |
| P2F01 | GCACAGACAATATTTTTGAATGGGGTCAGTA**GGTGTAGTTAATGGGAGT** |
| P2F02 | TTAACACCAGCACTAACAACTAATCGTTATTA |
| P2F03 | ATTTTAAAATCAAAATTATTTGCACGGATTCG |
| P2F04 | CCTGATTGCAATATATGTGAGTGATCAATAGT**GGTGTAGTTAATGGGAGT** |
| P2F05 | GAATTTATTTAATGGTTTGAAATATTCTTACC |
| P2F06 | AGTATAAAGTTCAGCTAATGCAGATGTCTTTC |
| P2F07 | CTTATCATTCCCGACTTGCGGGAGCCTAATTT |
| P2F08 | GCCAGTTAGAGGGTAATTGAGCGCTTTAAGAA**GGTGTAGTTAATGGGAGT** |

| | |
|---|---|
| P2F09 | AAGTAAGCAGACACCACGGAATAATATTGACG |
| P2F10 | GAAATTATTGCCTTTAGCGTCAGACCGGAACC |
| P2F11 | GCCTCCCTCAGAATGGAAAGCGCAGTAACAGT**GGTGTAGTTAATGGGAGT** |
| P2F12 | GCCCGTATCCGGAATAGGTGTATCAGCCCAAT |
| P2G01 | AGATTAGAGCCGTCAAAAAACAGAGGTGAGGCCTATTAGT**GGTGTAGTTAATGGGAGT** |
| P2G02 | XXXXXXXXXXXXXXXXATTCATTTTTGTTTGGATTATACTAAGAAACCACCAGAAG |
| P2G03 | XXXXXXXXXXXXXXXXCACCCTCAGAAACCATCGATAGCATTGAGCCATTTGGGAA |
| P2G04 | GTGATAAAAAGACGCTGAGAAGAGATAACCTTGCTTCTGTTCGGGAGA |
| P2G05 | GTTTATCAATATGCGTTATACAAACCGACCGT**GGTGTAGTTAATGGGAGT** |
| P2G06 | GCCTTAAACCAATCAATAATCGGCACGCGCCT |
| P2G07 | GAGAGATAGAGCGTCTTTCCAGAGGTTTTGAA |
| P2G08 | GTTTATTTTGTCACAATCTTACCGAAGCCCTTTAATATCA**GGTGTAGTTAATGGGAGT** |
| P2G09 | XXXXXXXXXXXXXXXXAACAATAACGTAAAACAGAAATAAAAATCCTTTGCCCGAA |
| P2G10 | XXXXXXXXXXXXXXXXAGCCACCACTGTAGCGCGTTTTCAAGGGAGGGAAGGTAAA |
| P2G11 | CAGGAGGTGGGGTCAGTGCCTTGAGTCTCTGAATTTACCGGGAACCAG |
| P2G12 | CCACCCTCATTTTCAGGGATAGCAACCGTACT**GGTGTAGTTAATGGGAGT** |
| P2H01 | CTTTAATGCGCGAACTGATAGCCCCACCAG**GGTGTAGTTAATGGGAGT** |
| P2H02 | CAGAAGATTAGATAATACATTTGTCGACAA |
| P2H03 | CTCGTATTAGAAATTGCGTAGATACAGTAC |
| P2H04 | CTTTTACAAAATCGTCGCTATTAGCGATAG**GGTGTAGTTAATGGGAGT** |
| P2H05 | CTTAGATTTAAGGCGTTAAATAAAGCCTGT |
| P2H06 | TTAGTATCACAATAGATAAGTCCACGAGCA |
| P2H07 | TGTAGAAATCAAGATTAGTTGCTCTTACCA |
| P2H08 | ACGCTAACACCCACAAGAATTGAAAATAGC**GGTGTAGTTAATGGGAGT** |
| P2H09 | AATAGCTATCAATAGAAAATTCAACATTCA |
| P2H10 | ACCGATTGTCGGCATTTTCGGTCATAATCA |
| P2H11 | AAATCACCTTCCAGTAAGCGTCAGTAATAA**GGTGTAGTTAATGGGAGT** |
| P2H12 | GTTTTAACTTAGTACCGCCACCCAGAGCCA |

Table S2: **Staples for each line-actant design to achieve line-accumulation**: Selected staples that feature an added overhang as an anchor-point in each variant of our line-actants.

| 6× | | 12× | | 24× | |
|---|---|---|---|---|---|
| dC | sT | dC | sT | dC | sT |
| P1A5 | P2C5 | P1A1 | P2C1 | P1A1 | P2C1 |
| P1A8 | P2C8 | P1A5 | P2C5 | P1A5 | P2C5 |
| P1C5 | P2E1 | P1A8 | P2C8 | P1A8 | P2C8 |
| P1C8 | P2E5 | P1A12 | P2C12 | P1A12 | P2C12 |
| P1E5 | P2G5 | P1C1 | P2E1 | P1B1 | P2D1 |
| P1E8 | P2G8 | P1C5 | P2E5 | P1B4 | P2D4 |
| | | P1C8 | P2E8 | P1B8 | P2D8 |
| | | P1C12 | P2E12 | P1B11 | P2D11 |
| | | P1E1 | P2G1 | P1C1 | P2E1 |
| | | P1E5 | P2G5 | P1C5 | P2E5 |
| | | P1E8 | P2G8 | P1C8 | P2E8 |
| | | P1E12 | P2G12 | P1C12 | P2E12 |
| | | | | P1D1 | P2F1 |
| | | | | P1D4 | P2F4 |
| | | | | P1D8 | P2F8 |
| | | | | P1D11 | P2F11 |
| | | | | P1E1 | P2G1 |
| | | | | P1E5 | P2G5 |
| | | | | P1E8 | P2G8 |
| | | | | P1E12 | P2G12 |
| | | | | P1F1 | P2H1 |
| | | | | P1F4 | P2H4 |
| | | | | P1F8 | P2H8 |
| | | | | P1F11 | P2H11 |

Table S3: **Anchor modules and oligonucleotides that comprise them.** Each construct is self-assembled through a quenching temperature ramp as discussed in the Methods at a final concentration of 2 $\mu$M and then diluted for GUV functionalisation. Constructs require strands in 1:1 stochiometric ratios.

| **Construct** | **Strands** |
|---|---|
| $\text{Anchor}_{\text{dC}}$ | $C_{\text{bb,chol}}$ + $C_{\text{b,chol}}$ |
| $\text{Anchor}_{\text{sT}}$ | $E_{\text{bb}}$ + $E_{\text{b,toc}}$ |

Table S4: **Sequences of oligonucleotide strands.** Abbreviations: TEG: triethyleneglycol.

| **Strand** | **Sequence (5' → 3')** |
|---|---|
| $C_{\text{b,chol}}$ | /Cholesteryl-TEG/CAATCACACCACAAACACCCAACACAACAACAAACC |
| $C_{\text{bb,chol}}$ | GTGTTTGTGGTGTGATTG/Cholesterol-TEG/ |
| $E_{\text{bb}}$ | CGCACTCTCACGCTAAAC |
| $E_{\text{b,toc}}$ | /Octyl-Tocopherol/GTTTAGCGTGAGAGTGCGACTCCCATTAACTACACC |
| $\text{Lin}^{f,*}$ | GCTCTCTCATCACTAC/AlexaFluor488/ |
| Extender | GTGTTGAGTAGTGAGATGGTAGTGATGAGAGAGCGTAGTGATGAGAGAGCGTAGT GATGAGAGAGC |

Table S5: **Sequences of domains added to staples for fluorescent beacon hybridisation.** Docking domains were added replacing the domains marked as XXXXXXXXXXXXXXXXX. Note that the extended beacon requires a stochiometric mixture of Extender + (3×)$Lin^{f,*}$.

| **Strand** | **Sequence (5' → 3')** |
|---|---|
| $Docking_{Lin^f}$ | GTAGTGATGAGAGAGC |
| $Docking_{Extender}$ | CATCTCACTACTCAACAC |

Table S6: **Parameters and values for numerical model.** The table lists the parameters that have been used to infer the density of tiles at the line from the experimental values of the line–partitioning $K^{\times n}_{\mathrm{p,int}}$ ($n$ = 6, 12, and 24). See Supplementary Discussion III for the definitions of the parameters.

| **Tiles** | $K^{n\times}_{\mathrm{p,int}}$ | $\delta K^{n\times}_{\mathrm{p,int}}$ | $\beta\Delta G_{\mathrm{p},L_{\mathrm{d}}}$ | $\beta\Delta G_{\mathrm{p},L_{\mathrm{o}}}$ | $\beta\Delta\tilde{G}_{\mathrm{p},L_{\mathrm{o}}}$ | $\beta\Delta G_{\mathrm{p,int}}$ | $\beta\Delta\tilde{G}_{\mathrm{p,int}}$ | $\delta$ [nm] | $L_1$ [nm] | $L_2$ [nm] | $\Delta\theta$ [deg°] |
|---|---|---|---|---|---|---|---|---|---|---|---|
| 6× | 79.9 | 44.7 | 0 | 6.6 | 5.91 | -4.8 | -4.15 | 38 | 75 | 80 | 101.3 |
| 12× | 44.9 | 25.9 | 0 | 13.2 | 11.26 | -9.6 | -7.58 | 38 | 75 | 80 | 37.5 |
| 24× | 52.4 | 29.4 | 0 | 26.4 | 21.42 | -19.2 | -15.68 | 33 | 75 | 80 | 32.6 |

Table S7: **Staple incorporation probabilities.** The table lists the Staple incorporation probabilities ($p$) used to compute Eq. S17. The values of $p$ were reported by Jungmann and co-workers[17] (see Supplementary Figure 14(b) in reference[17] )

| dC | | sT | |
|---|---|---|---|
| Staple | $p$ | Staple | $p$ |
| P1A12 | 0.699 | P2C12 | 0.823 |
| P1A8 | 0.914 | P2C8 | 0.855 |
| P1A5 | 0.85 | P2C5 | 0.903 |
| P1A11 | 0.602 | P2C1 | 0.704 |
| P1B11 | 0.833 | P2D11 | 0.914 |
| P1B8 | 0.85 | P2D8 | 0.876 |
| P1B4 | 0.769 | P2D4 | 0.855 |
| P1B1 | 0.774 | P2D1 | 0.79 |
| P1C12 | 0.737 | P2E12 | 0.833 |
| P1C8 | 0.821 | P2E8 | 0.952 |
| P1C5 | 0.79 | P2E5 | 0.946 |
| P1C1 | 0.667 | P2E1 | 0.785 |
| P1D11 | 0.882 | P2F11 | 0.812 |
| P1D8 | 0.941 | P2F8 | 0.828 |
| P1D4 | 0.871 | P2F4 | 0.86 |
| P1D1 | 0.871 | P2F1 | 0.715 |
| P1E12 | 0.753 | P2G12 | 0.764 |
| P1E8 | 0.925 | P2G8 | 0.823 |
| P1E5 | 0.882 | P2G5 | 0.812 |
| P1E1 | 0.839 | P2G1 | 0.715 |
| P1F11 | 0.86 | P2H11 | 0.871 |
| P1F8 | 0.893 | P2H8 | 0.785 |
| P1F4 | 0.833 | P2H4 | 0.823 |
| P1F1 | 0.742 | P2H1 | 0.479 |

Table S8: **Average probability of staple incorporation $p$ for dC and sT sets of anchor-points.** Average probabilities are computed from values in Table S7 for each line-actant design (see Table S4). In all cases, average probabilities for either set of anchor-points are $p \sim 0.8$, suggesting that an unbalance between anchor-modules on the tiles is unlikely.

| **Tile design** | 6× | 12× | 24× |
|---|---|---|---|
| Average $p$ for dC | 0.863 | 0.795 | 0.817 |
| Average $p$ for sT | 0.854 | 0.826 | 0.813 |

Table S9: **Sequence of toehold domain and Fuel strand.** Toehold domains were added at the 3' termini of dC staples $P1XX_{TH}$

.

| **Strand** | **Sequence (5' → 3')** |
|---|---|
| Toehold ($\alpha$) | GTTGTA |
| Fuel | TACAACCCAACACAACAACAAACC |
